# Longevity in *C. elegans* Eat mutants is largely attributable to reduced bacterial infection

**DOI:** 10.64898/2026.03.11.711062

**Authors:** Hongyuan Wang, Yuan Zhao, Faria Athar, Jennifer N. Lohr, Zhou Yang, Bruce Zhang, Hannah Chapman, Ioan Marcu, Mila Penzel, David Gems

## Abstract

Dietary restriction (DR) extends lifespan in many animal species. In *C. elegans*, Eat mutants with pharyngeal defects that impair feeding exhibit reduced growth rate and fertility and are typically long-lived, suggesting a DR effect. We report that Eat mutant longevity is largely or wholly a consequence of suppression of feeding activity-dependent infection of the pharynx by their *E. coli* food source. *eat-2* mutants, widely used as a DR model, were among only 2/8 Eat mutants tested whose longevity were to any degree independent of bacterial infection. Moreover, among Eat mutants, phenotypic indicators of reduced nutrition correlated with one another, yet not with longevity. These findings document how, if infection is excluded, Eat mutants experience reduced nutrition, but in most cases not longevity, i.e. life-extending DR effects are not typical of Eat mutants. Thus, *eat-2* longevity is partially due to infection resistance rather than DR, and residual, pharyngeal infection-independent longevity (contributing ∼40% of the total increase in lifespan) could reflect DR, or alternatively some other consequence of their nicotinic acetylcholine receptor defect.

## Introduction

Dietary restriction (DR, sometimes referred to as caloric restriction), the controlled reduction of food intake below that which is optimal for growth and reproduction, is an experimental intervention that extends lifespan in laboratory model organisms ranging from budding yeast to rodents, sometimes markedly (reviewed in (Schmauck-Medina et al., 2026)). The term DR is often used to refer to food restriction that reduces nutrition and extends lifespan, rather than to reduced nutrition per se. To avoid confusion in this article we will distinguish DR (reduced nutrition that extends lifespan) from reduced nutrition as such. While there is a paucity of clear evidence of anti-aging DR effects in primates and humans, as opposed to mere rescue from the life-shortening effects of over-eating and obesity (Mattison et al., 2017; Most et al., 2017; Pifferi et al., 2018), DR in model organisms is clearly useful at least for investigating mechanisms of aging.

The tiny, free-living nematode *Caenorhabditis elegans* is a convenient animal model for studying the biology of aging, thanks in particular to its short lifespan and the identification of many genes mutation of which can increase its lifespan (the Age phenotype). It has also been used extensively to investigate the biology of DR (Kapahi et al., 2016). As many as 10 different approaches have been used to subject *C. elegans* to reduced nutrition (Greer and Brunet, 2009; Kapahi et al., 2016). Among these, perhaps the most reliable and versatile is bacterial DR (bDR), which involves controlled dilution of *E. coli* in liquid (S medium), wherein its proliferation is prevented by the absence of nutrients (Klass, 1977).

Another, frequently used approach is employment of *eat* (EATing abnormal) mutants with pharyngeal defects that interfere with normal ingestion of food. Here, of a number of *eat* mutants originally shown to be long lived (Lakowski and Hekimi, 1998), *eat-2* mutants have been used almost exclusively, as a DR model in numerous reports, of which 77 examples are listed in Supplementary Table 1. Eat mutants show signs of reduced nutrition (e.g. slow growth, reduced fertility and a pale, malnourished appearance) and, importantly, are typically long-lived. This supports the view that Eat mutant longevity, including that of *eat-2*, is a consequence of reduced nutrition, i.e. that they experience DR (Lakowski and Hekimi, 1998).

However, it remains possible that this interpretation is not correct. A major difference between the classic rodent DR model and *C. elegans* is that in the latter the food source (the bacterium *Escherichia coli*) is also a life-shortening pathogen. Standard *C. elegans* culture conditions involve maintenance on an agar plate with a lawn of live *E. coli* (usually the strain OP50) (Brenner, 1974). The *E. coli* colonizes, invades and, in later life, kills *C. elegans*, and prevention of this by blocking bacterial proliferation increases lifespan substantially (Garigan et al., 2002; Gems and Riddle, 2000; Podshivalova et al., 2017; Zhao et al., 2017). Because the *C. elegans* laboratory food source is also a pathogen, it can be difficult to distinguish whether effects on lifespan of reducing food levels are attributable to reduced nutrition or reduced infection (Walker et al., 2005).

In a previous study we described how in wild-type *C. elegans* a subset (∼40%) of aging hermaphrodites die relatively early with an *E. coli* infection wherein the pharynx becomes enlarged and swollen (P [“big P”] death) (Zhao et al., 2017) (Fig. 1A). Individuals that escape P death, and therefore live longer, subsequently die with an atrophied pharynx (p [“small p”] death), but also from *E. coli* infection (since prevention of *E. coli* proliferation increases p lifespan).

**Figure 1.**
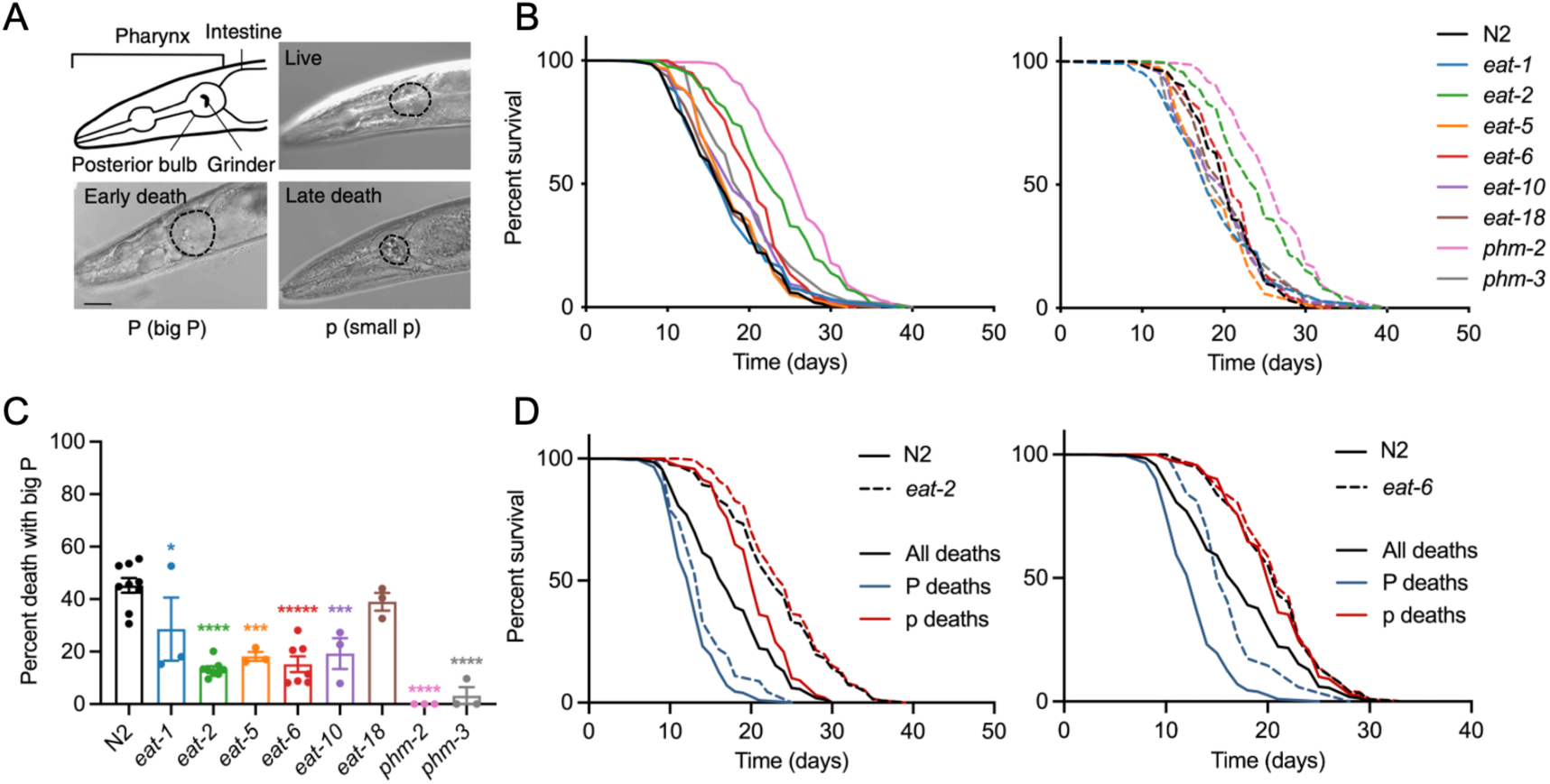
Eat mutants longevity is largely attributable to reduced P death frequency. (**A**) Nematode corpses with an enlarged or an atrophied pharynx (P and p deaths, respectively) (Zhao et al., 2017). For comparison, the pharynx of a healthy 10-day-old adult worm is shown (posterior bulb outlined). Scale bar, 40 μm. (**B**) Lifespans of the whole population (left) and p sub-population (right) of Eat mutants on proliferating *E. coli* OP50. *N* ≥3 trials. See Supplementary Table 2 for statistics. (**C**) P death frequency of Eat mutants on proliferating *E. coli* OP50. *N* >3 trials. (**D**) Deconvolved survival curves for N2 *eat-2* and *eat-6*.

Treatments that alter lifespan can differentially affect the two types of death, in terms of their frequency and timing. This can be assessed by combining mortality and necropsy data, and examining P and p sub-population survival separately (mortality deconvolution). Notably, an earlier, limited analysis of *eat-2(ad1116)* established that its longevity was caused largely by a reduction in P death frequency, apparently due to reduced mechanical injury (mechanical senescence) of the cuticular lining of the pharynx, resulting from the diminished pharyngeal pumping rate, rather than to reduced nutrition and DR (Zhao et al., 2017).

Clearly, for studies of DR in *C. elegans*, it is important to establish whether or not the widely used putative DR model, *eat-2* mutants, are in fact long-lived due to reduced nutrition. In this study we assess this more fully by reexamining aging in a range of Eat mutants, using mortality deconvolution analysis along with other approaches. We also assess which Eat mutant traits are correlated with mortality parameters (overall lifespan, P and p lifespan, and P and p frequency), including pharyngeal infection in early adulthood, reduced nutrition-linked parameters (including retarded development and reduced fertility), and bacterial lawn avoidance (Kumar et al., 2019). Our findings demonstrate that Eat mutant longevity is mainly attributable to reduced pharyngeal infection, and demonstrate that *eat-2* mutant longevity is at least partly, and possibly even wholly independent of reduced nutrition.

## Results

### *eat-2* and *phm-2* reduce P death and induce p Age

To reassess the causes of Eat mutant longevity, we studied a panel of 8 Eat mutants (detailed in Table 1). Of these, 6 were included in the initial *eat* mutant longevity study: *eat-1(ad427)*, *eat-2(ad1116)*, *eat-5(ad464)*, *eat-6(ad467)*, *eat-10(ad606)*, and *eat-18(ad1110)*, of which all but *eat-5* were previously found to extend lifespan (Lakowski and Hekimi, 1998). Also included was *phm-2(ad597),* which causes a pharyngeal defect that allows unmasticated, live bacteria to readily enter the intestinal lumen, and also extends lifespan (Kumar et al., 2019). Phm (PHaryngeal Muscle) mutants exhibit defects in pharyngeal muscle, identifiable as altered muscle birefringence (Avery, 1993). *phm-3(ad493)* was also included for comparison with *phm-2*.

**Table 1.** Overview of Eat mutant genes used in this study.

| Mutation | Gene function | Feeding phenotype(Avery, 1993) | Previously observed effect on mean lifespan | Original source for effect on lifespan |
| --- | --- | --- | --- | --- |
| *eat-1(ad427)* | Unknown | Slow, irregular feeding | +33% | (Lakowski and Hekimi, 1998) |
| *eat-2(ad1116)* | Synaptic transmission (nicotinic acetylcholine receptor subunit) | Slow, regular feeding | +57% | (Lakowski and Hekimi, 1998) |
| *eat-5(ad464)* | Gap junction hemichannel | Unsynchronized contraction | -6% | (Lakowski and Hekimi, 1998) |
| *eat-6(ad467)* | Na<sup>+</sup>/K<sup>+</sup> exchange | Relaxation-deficient | +37% | (Lakowski and Hekimi, 1998) |
| *eat-10(ad606)* | Unknown | Slippery pharynx | +8% | (Lakowski and Hekimi, 1998) |
| *eat-18(ad1110)* | Synaptic transmission (acetylcholine-gated channel complex subunit) | Slow, regular feeding | +15% | (Lakowski and Hekimi, 1998) |
| *phm-2(ad597)* | Unknown function, SAFB (human protein scaffold attachment factor B) homolog | Abnormal grinder position | +33% | (Kumar et al., 2019) |
| *phm-3(ad493)* | Unknown | Feeble contraction | ND | ND |

As a first step, all mutant strains were checked for the presence of the background mutation *fln-2(ot611)*, which reduces P death frequency and therefore early mortality, and suppresses *eat-2* longevity (Chang et al., 2026; Zhao et al., 2019) (since *eat-2* mutant longevity is largely attributable to reduction of P death frequency). Two strains proved to contain *ot611*, DA531 *eat-1(ad427) IV; fln-2(ot611) X*, and DA493 *phm-3(ad493) III; fln-2(ot611) X*. The mutants were backcrossed with N2 to replace *fln-2(ot611)* with *fln-2(+)*, to generate strains GA6005 *eat-1(ad427)* and GA6003 *phm-3(ad493)*.

We then assessed whether, across the Eat mutant panel, longevity is attributable to reduced P frequency, as shown previously in *eat-2* mutants. To this end all strains were first subjected to survival, necropsy and mortality deconvolution analysis. Of the 8 mutants tested, 5 (*eat-2, eat-6, eat-10, phm-2* and *phm-3*) showed a significant increase in overall mean lifespan, by +35.10%, +20.73%, +9.98%, +54.16% and +15.24%, respectively (*p* < 0.0001*, p* < 0.0001, *p* = 0.0015, *p* < 0.0001 and *p* = 0.0001, respectively, log rank test; summed data from *N* ≥ 3; Fig. 1B left, Supplementary Table 2). For raw data for all survival trials, see Supplemental Dataset 1. Here our results largely reproduce earlier findings (Lakowski and Hekimi, 1998), with the exception that *eat-1(ad427)* and *eat-18(ad1110)* did not extend lifespan in our hands, for reasons unknown.

Next we deconvolved the Eat mutant mortality data into its P and p death components. Reductions in P death frequency were observed with *eat-2*, *eat-5*, *eat-10*, *phm-2* and *phm-3*, but not *eat-18*, much as previously reported (Zhao et al., 2017); *eat-1* and *eat-6* also reduced P death frequency (Fig. 1C).

Notably, mean p lifespan was extended in only 2 out of the 8 mutants, in *eat-2* (+18.01%) and *phm-2* (+27.38%) (*p* < 0.0001 in both cases; Fig. 1B right, Supplementary Table 2); in our previous study a non-significant trend to increased p lifespan was seen with *eat-2(ad1116)* (*p* = 0.081) (Zhao et al., 2017). Thus, in *eat-6*, *eat-10* and *phm-3* mutants, longevity is solely attributable to reduced P death frequency.

That lifespan can be increased in Eat mutants solely by reduction in P death frequency is also evident from deconvolved survival curves, showing effects on P and p subpopulations. For example, while *eat-2* modestly increases p lifespan, *eat-6* does not (Fig. 1D; for deconvolved survival curves for other mutants, see Fig. S1).

Earlier evidence suggests that the high, wild-type rate of pharyngeal pumping in early adulthood promotes bacterial infection by causing mechanical injury and perforation of the pharyngeal cuticle (Zhao et al., 2017). Consistent with this, across Eat mutants P death frequency was positively correlated with the severity of pharyngeal infection earlier in life, showing a trend on D4 (day 4 of adulthood; *p* = 0.053) and a significant correlation on D8 (*p* = 0.0024) (Fig. 2A,B). Moreover, overall lifespan was also negatively correlated with the severity of pharyngeal infection earlier in life, showing a trend on D4 (*p* = 0.083) and a significant correlation on D8 (*p* = 0.034) (Fig. 2C). These findings support the view that in Eat mutants reduced pharyngeal pumping rates in young adults prevent early life *E. coli* invasion of the pharynx, that in later life develops into life-shortening, lethal infection.

**Figure 2.**
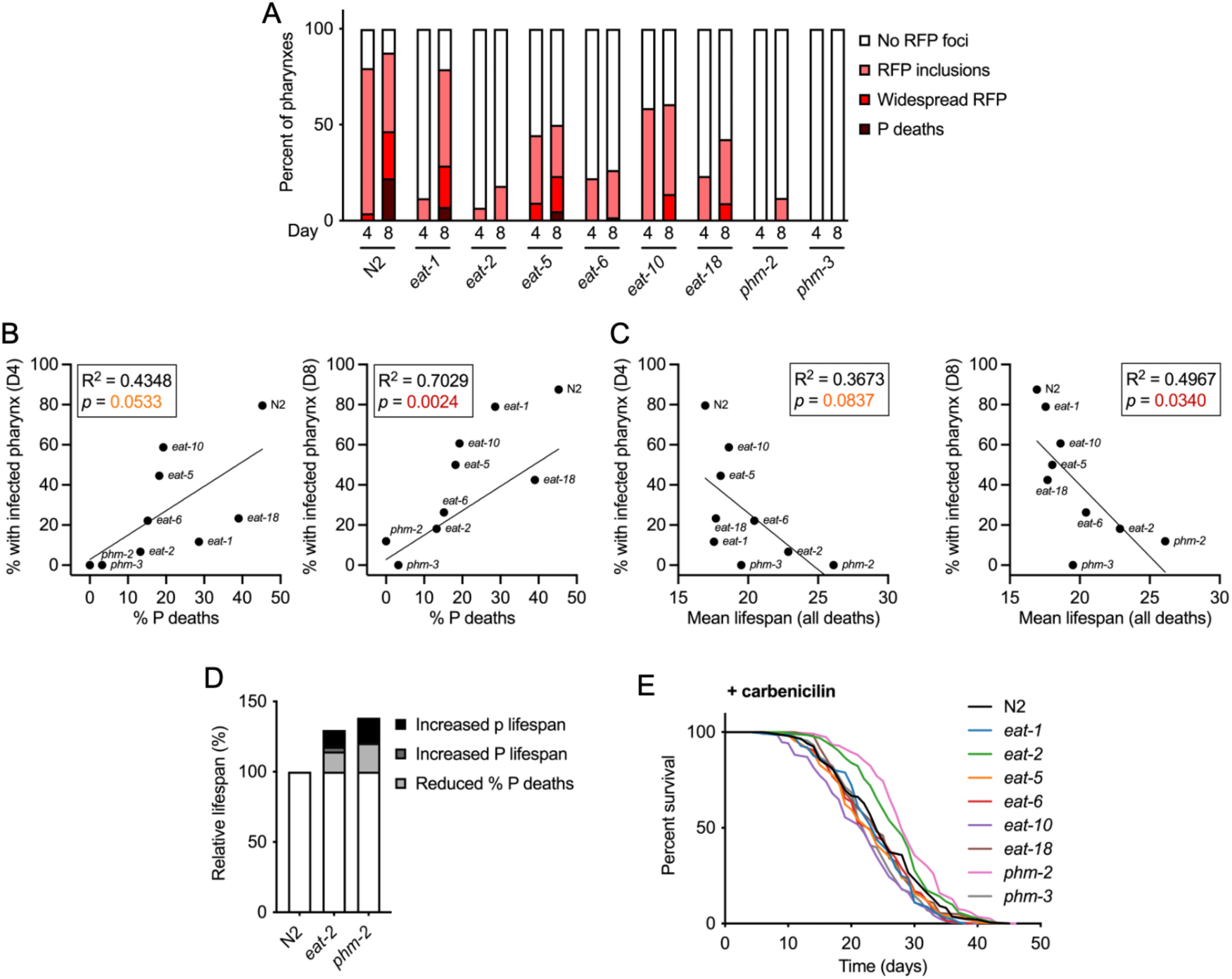
Eat mutants longevity is largely attributable to reduced P death frequency. (**A**) Pharyngeal invasion by OP50-RFP on day 4 and day 8 of adulthood. *n* > 10 per time point. (**B**) Regression analysis of early pharyngeal infection and P death frequency, day 4 (left) and day 8 (right). (**C**) Regression analysis of early pharyngeal infection and overall lifespan, day 4 (left) and day 8 (right). (**D**) Respective contributions to changes in overall lifespan of reduction in P death frequency and p Age (extended lifespan of p subpopulation). (**E**) Lifespans of Eat mutants on carbenicillin-treated (i.e. non-proliferating) *E. coli*. *N* = 4 trials. See Supplementary Table 3 for statistics.

*E. coli* OP50-RFP was also used to investigate the origins and route of entry of pharyngeal infection. The earliest signs of infection, from day 2 of adulthood onward, were often evident as small red fluorescence puncta within pharyngeal tissue near to the grinder (Fig. S2A,B). These appeared at a significantly higher frequency in the anterior hemisphere of the terminal pharyngeal bulb (Fig. S2C), suggesting a possible access route for infection from the anterior pharyngeal lumen into the tissue. The bias towards greater infection in the anterior of the pharyngeal bulb decreased with increasing level of infection (Fig. S2D) and with increasing age (Fig. S2E).

Focusing on the two mutants exhibiting p Age (extended longevity in the p sub-population), we next estimated the relative impact of reduced P vs p Age in overall *eat-2* and *phm-2* Age. This was achieved by calculating the predicted effect on mean wild-type lifespan of each individual change (i.e. in %P vs p Age). For *eat-2(ad1116)*, reduced P death frequency, and p longevity caused 14.41% and 11.87% increases, respectively; for *phm-2(ad597)*, the respective values were 20.39% and 18.05% (Fig. 2D). Thus, reduced P death frequency contributes somewhat more than p Age to the overall Age phenotype in both mutants. Moreover, p Age appears to be a slightly greater contributor to overall Age in *phm-2* than *eat-2* mutants (46.96% vs 40.03%, respectively).

### *eat-2* and *phm-2* p Age is not attributable to infection resistance

Blocking bacterial proliferation extends *C. elegans* lifespan by preventing P death and extending p lifespan (Zhao et al., 2017). To test the effects of the 8 Eat mutations on lifespan without the confounding effects of bacterial infection, the antibiotic carbenicillin was used. Here again only *eat-2* and *phm-2* extended lifespan (+10.98%, *p* = 0.0078, +17.21%, *p* < 0.0001, respectively; Fig. 2E, Supplementary Table 3). This provides further evidence that *eat-6*, *eat-10* and *phm-3* mutant longevity is not attributable to reduced nutrition, but rather to resistance to pharyngeal infection resistance.

By contrast, the fact that *eat-2* and *phm-2* Age occur on non-proliferating *E. coli* suggests that *eat-2* and *phm-2* p Age (no carbenicillin) are not attributable to resistance to bacterial infection. Here our findings are consistent with relatively modest increases in *eat-2* mutant lifespan described in previous studies using *E. coli* that was UV-irradiated (Calvert et al., 2016; Kaeberlein et al., 2006; Kumar et al., 2019) (+16%, +17%, +10%, respectively), or heat killed (+5.9%)(Statzer et al., 2022), and diminished *eat-2* life extension when maintained on less pathogenic bacteria, *Comamonas sp*. (Kauffman et al., 2010) or *Bacillus subtilis* (Sanchez-Blanco and Kim, 2011), relative to populations on *E. coli*.

One possibility is that bacterial proliferation causes reduced nutrition in Eat mutants, such that carbenicillin increases nutrition, thereby suppressing a DR effect. To test this we directly compared brood size and reproductive schedule in the 8 Eat mutants in the absence and presence of carbenicillin, but no evidence of rescue of reduced fertility on live *E. coli* was detected (Fig. S3). Thus, it is unlikely that carbenicillin suppresses a DR effect.

To investigate whether the effects of *eat-2(ad1116)* are typical of *eat-2* mutants, two further mutants were characterized, DA465 *eat-2(ad465)* and DA1113 *eat-2(ad1113)*. First, it was established that both strains carry the *fln-2(+)* allele. Next, lifespan and necropsy analysis was performed on proliferating *E. coli*. While all three alleles increased overall lifespan, the effect of *ad1116* appeared slightly greater than that of *ad465* and *ad1113* (+34.8%, +20.8% and +23.1%, respectively) (*N* = 2, Fig. S4A, Supplementary Table 4). The three alleles caused similar reductions in P death frequency (Fig. S4B). Notably, while *ad1116* significantly increased p lifespan (+13.17%, *p* = 0.0002), as previously seen (Fig. 1B right, 1D left), *ad465* and *ad1113* did not (Fig. S4C, Supplementary Table 4).

Altogether, these results show that *eat-2* is atypical among Eat mutants, which do not usually increase lifespan in the absence of bacterial proliferation. Importantly, this greatly weakens the initial claim that *eat-2* Age is attributable to reduced nutrition, i.e. reflects DR. To emphasize: this claim was based on the observation that Eat mutants *in general* are both malnourished and long lived (Lakowski and Hekimi, 1998). This in turn raises the possibility that *eat-2* p Age is not a consequence of the Eat phenotype, but rather an unrelated pleiotropic effect arising from the *eat-2* mutant defect in cholinergic signaling (McKay et al., 2004).

### Indicators of reduced nutrition are associated with reduced P death but not increased lifespan

One plausible reason for thinking that Eat mutant longevity reflects DR is the strong evidence that they experience reduced nutrition, including their scrawny, starved appearance, delayed development, and reduced fertility (Avery, 1993; Lakowski and Hekimi, 1998). That only *eat-2* and *phm-2* mutants exhibit infection-independent longevity suggests the hypothesis that they experience a greater reduction in nutrition than the other Eat mutants. To investigate this, we compared several metrics of reduced nutrition in the 8 mutants: developmental delay, reduced adult body size, and reduced and delayed progeny production.

All 8 Eat mutants showed some degree of developmental delay, with the strongest effects seen in *eat-1* and *eat-6* (Fig. 3A). Larval body size, measured 48 hr after egg lay, was reduced in all *eat* mutants (Fig. 3B). Overall brood size was significantly reduced in all cases, and most strongly for *eat-1* (Fig. 3C). In all mutants a delay in reproductive schedule was seen, evident as an increase in later (day 5) progeny production (Fig. 3D,E).

**Figure 3.**
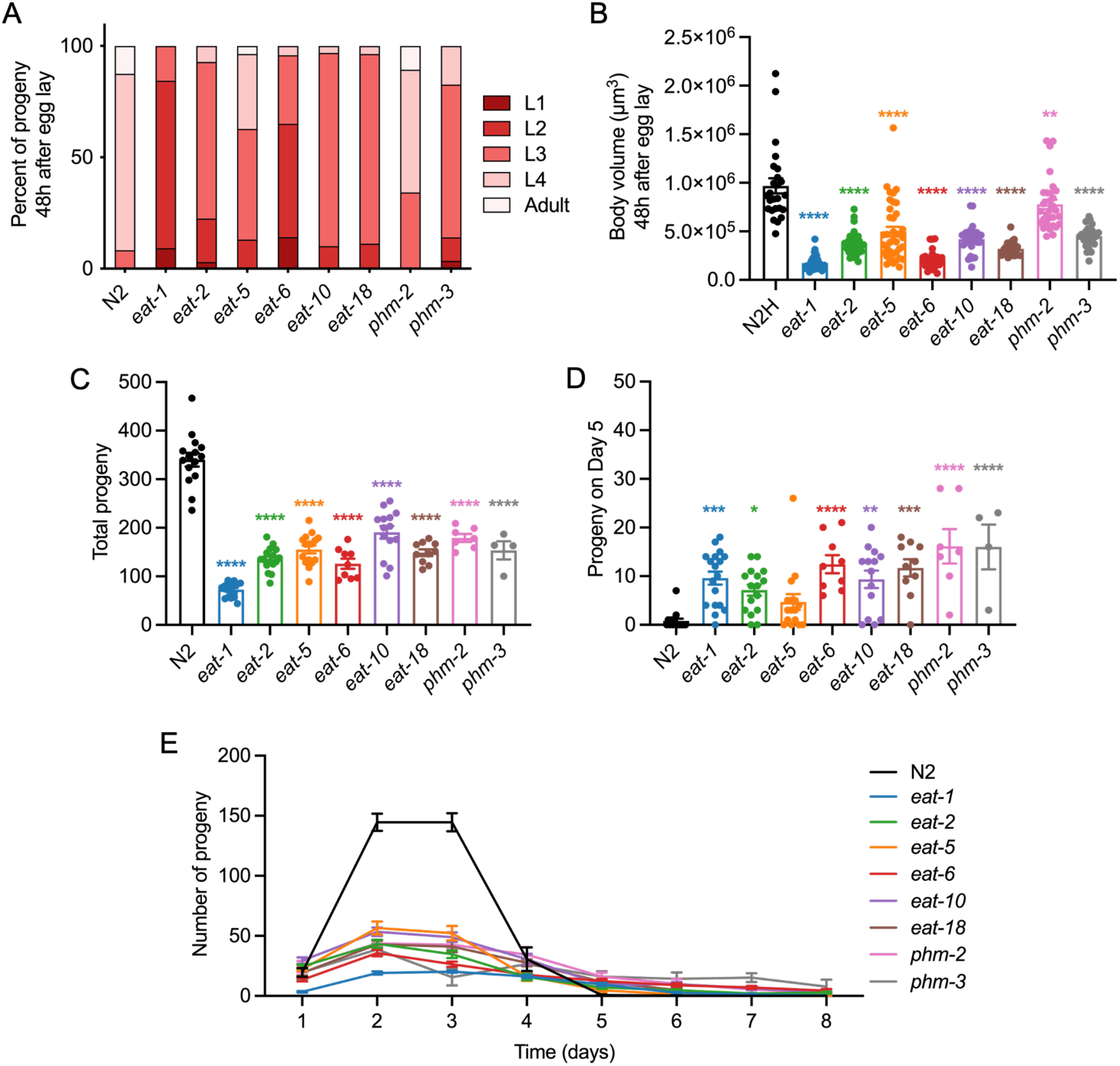
Traits indicative of reduced nutrition in Eat mutants. (**A**) Delayed development. (**B**) Reduced body size (48 hr). (**C**) Reduced brood size. (**D**,**E**) Delayed fertility schedule. (**D**) Increased progeny on day 5 of adulthood, due to delayed fertility schedule. (**D**) Overall fertility schedules, showing reduced early fertility and increased late fertility. All trials were performed at 20°C on proliferating *E. coli*.

We then examined correlations between metrics of reduced nutrition. As one would expect, a strong positive correlation was seen between brood size and body size at both 48 hr and 96 hr after egg laying (R^2^ = 0.77, *p* = 0.0017, and R^2^ = 0.81, *p* = 0.0009, respectively; Fig. S5A). Brood size and body size both showed some positive correlations with incidence of pharyngeal infection on D4 but not D8 of adulthood (Fig. S5B-D). This is in line with the view that reduced pharyngeal pumping rate reduces growth, fertility and pharyngeal infection.

Next we assessed whether severity of reduced nutrition-associated metrics are predictive of the degree of lifespan extension, as one would expect if Eat mutant longevity is attributable to DR. Regression analysis of developmental delay, adult body size, brood size and reproductive delay against lifespans of total populations, or p subpopulations, or carbenicillin-treated populations was performed (Fig. 4A-H, Fig. S6A-D). Strikingly, no significant correlations were detected with any of the 3 lifespan metrics. Given the absence of a general correspondence between metrics of reduced nutrition and longevity phenotypes among Eat mutants, the claim that *eat-2* and *phm-2* longevity is a consequence of DR is somewhat tenuous. However, it remains possible that an underlying relationship between reduced nutrition-associated traits and lifespan is obscured by pleiotropic effects of individual mutations on lifespan.

**Figure 4.**
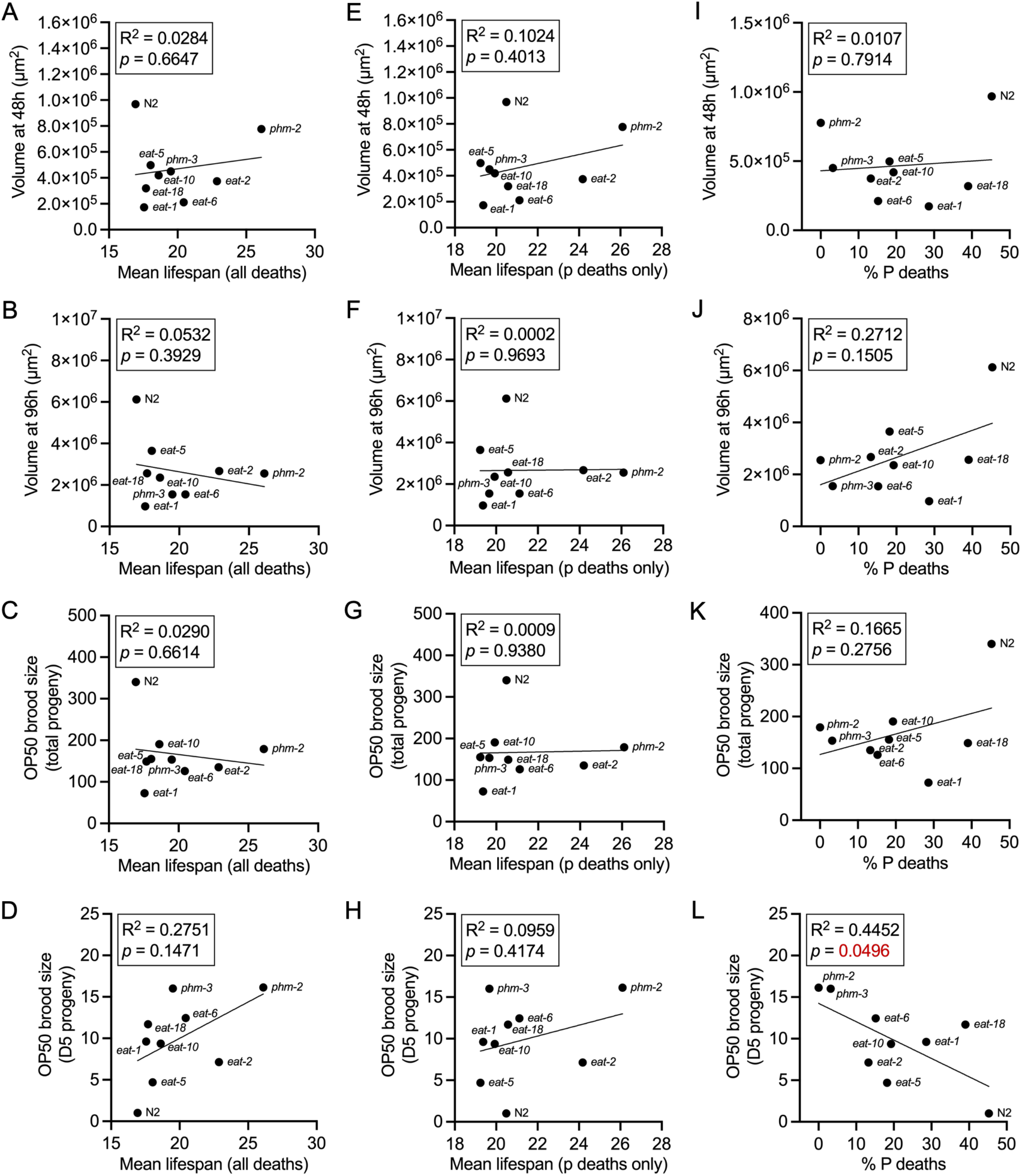
Correlation between metrics of reduced nutrition and aging in Eat mutants. (**A-D**) Correlations with overall lifespan. (**A**) With developmental delay. (**B**) With day 1 body size (estimated volume). (**C**) With brood size. (**D**) With day 5 progeny count (reproductive delay). (**E-H**) Correlations with p lifespan. (**E**) With developmental delay. (**F**) With day 1 body size (estimated volume). (**G**) With brood size. (**H**) With day 5 progeny count (reproductive delay). (**I-L**) Correlations with % P deaths. (**I**) With developmental delay. (**J**) With day 1 body size (estimated volume). (**K**) With brood size. (**L**) With day 5 progeny count (reproductive delay).

We also noted, as expected, a significant negative correlation between overall lifespan and P death frequency (Fig. S6E) and, also as expected, a positive correlation between lifespans of p subpopulations (no carbenicillin) and carbenicillin-treated populations (Fig. S6F).

We had also expected to see negative correlations between metrics of reduced nutrition and P death frequency, given the hypothesis that reduced pharyngeal pumping rate reduces bacterial invasion of pharyngeal tissue. However, this was only detected in one instance: delayed fertility (D5) (Fig. 4I-L). This could imply that the effect of any given *eat* or *phm* mutation on mechanical injury to the pharynx (Zhao et al., 2017) is not a simple function of its effect on nutrition.

### *eat-2* and *phm-2* Age are not attributable to lawn avoidance

Mutation of *eat-2* and *phm-2* cause bacterial lawn avoidance behavior, which has been suggested to contribute to life extension (Kumar et al., 2019; Singh and Aballay, 2019). This effect may be attributable to an innate immune response caused by earlier entry of live bacteria into the intestine due to the pharyngeal defects (Kumar et al., 2019). To further probe the lawn avoidance DR hypothesis we first compared lawn avoidance in N2 and the 8 Eat mutants. Significant levels of lawn avoidance (i.e. increased proportions of animals observed off the bacterial lawn) were seen in *eat-2* and *phm-2* mutants, as previously reported (Kumar et al., 2019), and also in *eat-10* and *phm-3* mutants (Fig. 5A left).

**Figure 5.**
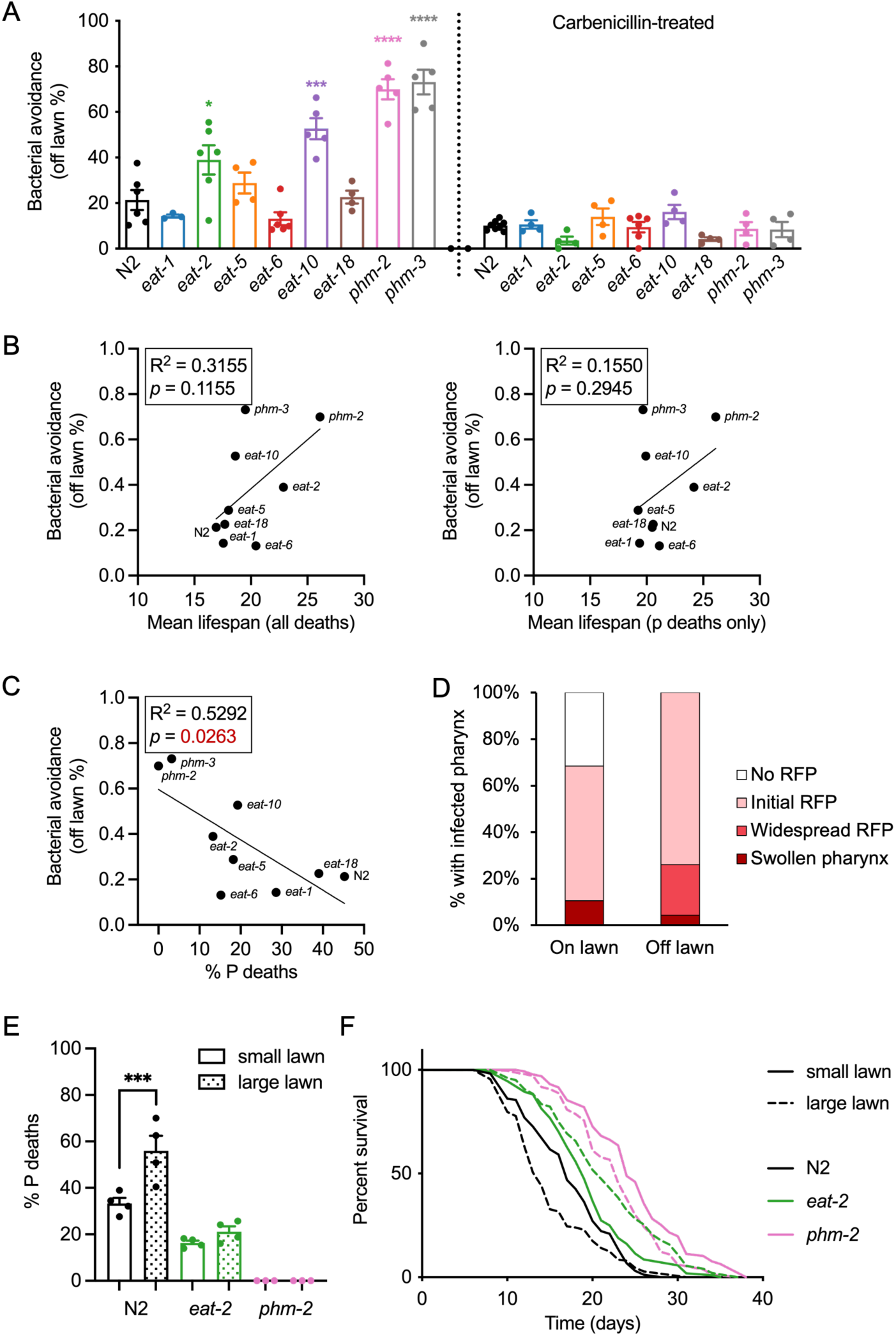
*eat-2* and *phm-2* Age are not attributable to lawn avoidance. (**A**) Many Eat mutants exhibit lawn avoidance (left), which is abrogated by preventing bacterial proliferation using carbenicillin (right). (**B**) Lawn avoidance does not correlate with total population lifespan (left) or p lifespan (right) among Eat mutants. (**C**) Lawn avoidance is strongly correlated with P death frequency among Eat mutants. (**D**) Increased pharyngeal infection in N2 hermaphrodites found outside the bacterial lawn (day 6 of adulthood). (**E**) Preventing lawn avoidance increases frequency of P death in N2 but not *eat-2* (*phm-*2 mutants do not exhibit P death). Mann-Whitney test, \*\*\**p*<0.001. (**F**) Abrogating lawn avoidance does not shorten lifespan in *eat-2* and *phm-2* mutants.

*eat-2* and *phm-2* lawn avoidance behavior appears to be dependent upon bacterial proliferation, since it was prevented by UV irradiation of *E. coli* (Kumar et al., 2019). Confirming this, blocking bacterial proliferation with carbenicillin prevented lawn avoidance in all cases (Fig. 5A right).

If lawn avoidance increases lifespan, one might expect a positive correlation between lawn avoidance and lifespan. However, we detected no correlation between lawn avoidance and either overall lifespan (Fig. 5B left) or p lifespan (Fig. 5B right). What we did observe was a significant negative correlation between lawn avoidance and P death frequency (*p* = 0.026, Fig. 5C). This and the fact that preventing bacterial proliferation reduces lawn avoidance (Fig. 5A) (Kumar et al., 2019) suggests that in mutants with lower P death frequency such as *phm-2* and *phm-3*, the reduction of pharyngeal function that suppresses P death also increases entry of live *E. coli* into the intestinal lumen, triggering a lawn avoidance response.

A further possibility is that pharyngeal infection induces lawn avoidance. To test this, frequency of pharyngeal infection was compared in N2 hermaphrodites found on or off the *E. coli* lawn on D6 of adulthood. The proportion of infected animals proved to be higher in those that were off the lawn (Fig. 5D), supporting this possibility.

We then asked: is lawn avoidance a simple function of degree of reduced nutrition? To this end we performed regression analyses of lawn avoidance frequency with two metrics of reduced nutrition: body size and brood size, but no correlations were detected (Fig. S7A-C), arguing against it being a consequence of DR.

Regression analysis of lawn avoidance frequency against infection levels on D4 and D8 of adulthood detected only a trend for a negative correlation on D8 (*p* = 0.069, Fig. S7D,E). This is likely attributable to the two Phm mutants, where greatly reduced mechanical senescence, preventing pharyngeal infection, is coupled to increased entry of intact *E. coli* into the intestinal lumen, triggering lawn avoidance.

To further test the role of lawn avoidance on *eat-2* and *phm-2* mutant longevity, avoidance was prevented by using agar plates where the bacterial lawn covers the entire plate (large lawns), as described (Kumar et al., 2019). Preventing lawn avoidance increased P death frequency in N2 but not *eat-2* or, unsurprisingly, *phm-2* which prevents P death altogether (Fig. 5E). Notably, though preventing lawn avoidance did significantly reduce N2 lifespan (−13.1%, *p* = 0.0054), it did not do so in *eat-2* and *phm-2* mutants, but modestly increased it in the former (+11.7%, *p* = 0.0016, Fig. 5F, Supplementary Table 5), an effect for which we have no ready explanation.

In the prior study in which prevention of lawn avoidance abrogated *phm-2* mutant longevity, high levels of death from internal hatching of larvae (matricide) were observed, and not censored from survival analysis (Kumar et al., 2019). In our data matricidal deaths were censored. However, all non-aging-related deaths were scored, and frequencies on small vs large lawns were in N2 9.4% vs 10.4%, in *eat-2* mutants 27.7% vs 31.0%, and in *phm-2* mutants 28.6% vs 60.8%, respectively (Supplementary Table 5); the increase on large lawns in *phm-2* mutants was largely attributable to matricide.

The increase in matricide frequency caused by maintaining *phm-2* mutants on large lawns was suppressed by addition of carbenicillin (Fig. S8A), implying that bacterial proliferation triggers matricide in this strain. Plausibly, this reflects the *phm-2* open pharynx phenotype that allows intact, unmasticated bacteria to enter the intestinal lumen (Avery, 1993; Darby et al., 2002; Labrousse et al., 2000). We also noted a reduction in body size in *phm-2* mutants on large lawns which, again, was suppressed by carbenicillin (Fig. S8B); thus, in the case of *phm-2* mutants on large lawns, proliferative *E. coli* can reduce growth, a metric of reduced nutrition. These results suggest that *phm-2* mutants avoid the lawn to try to reduce ingestion of unmasticated *E. coli*, and when they cannot do so are so afflicted that growth and egg laying is reduced, the latter perhaps reflecting a pathogen response (Mosser et al., 2011). In conclusion, these findings argue that *eat-2* and *phm-2* longevity are not the consequence of increased lawn avoidance.

## Discussion

*eat-2* mutants are widely used as a convenient genetic model for studying the biology of DR using *C. elegans* (Supplemental Table 1). At the time of writing (July 2026) the original 1998 article reporting that *eat-2* extends lifespan by causing DR (Lakowski and Hekimi, 1998) has been cited 466 times in PubMed. However, findings presented here identify several issues with the use of *eat-2* mutants as a DR model.

First, although Eat mutants are typically long-lived, in the majority of cases this is attributable solely to reduced frequency of death from pharyngeal *E. coli* infection (P death). When such infection is prevented, only 2/8 mutants tested, *eat-2(ad1116)* and *phm-2(ad597)*, were long-lived (Fig. 2E, Supplementary Table 3). The initial observation that Eat mutants as a category are typically long-lived strongly implied that *eat-2* longevity reflects DR (Lakowski and Hekimi, 1998); however, this critical implication no longer holds.

Second, only ∼40% of the *eat-2* longevity is not attributable to reduced P death frequency (Fig. 2D). Thus, even if infection-independent *eat-2* Age is attributable to reduced nutrition, the presence of two distinct mechanisms of life-extension clouds and confounds interpretation of experimental results.

Third, if Eat mutant longevity is attributable to reduced nutrition (as the DR interpretation argues) then, among Eat mutants, traits arising from reduced nutrition, such as delayed development, reduced body size and brood size, should show a positive correlation with longevity, yet they do not (Fig. 4A-H, Fig. S6A-D); notably *eat-1* and *eat-6* mutants exhibit stronger indications of reduced nutrition than *eat-2* or *phm-2* mutants (Fig. 3A-C). We also demonstrate that *eat-2* and *phm-2* longevity is not the result of DR caused by increased lawn avoidance (Fig. 5).

These findings support the view that reduced pharyngeal pumping rate in Eat mutants protects them from high pumping rate-dependent promotion of *E. coli* infection in the pharynx. This may reflect reduced mechanical senescence of the pharyngeal cuticle in early adulthood, when pumping rate is at its peak (Zhao et al., 2017). A further possibility, suggested previously, is that it protects against contraction-related injury (again, mechanical senescence) that otherwise causes sarcopenia in pharyngeal muscle (Chow et al., 2006).

### Are *eat-2* and *phm-2* p Age attributable to DR?

This remains uncertain, but it is a possibility. That *eat-2* and *phm-2* extend lifespan in the absence of proliferating bacteria indicates action of a mechanism independent of infection resistance, that could therefore be a consequence of reduced nutrition. Alternatively, however, it could reflect pleiotropic effects of these two mutations, unconnected to reduced nutrition. *eat-2* encodes a subunit of a nicotinic acetylcholine receptor that acts in pharyngeal muscle (McKay et al., 2004); possibly other effects of deficiency of this receptor on signaling or physiology extend lifespan.

*phm-2* encodes a protein of unknown function homologous to the human protein scaffold attachment factor B (Kumar et al., 2019). *phm-2(ad597)* caused the greatest increase in overall lifespan in the mutants examined, with or without infective bacteria. The severity of the feeding defect, wherein the masticatory function of the pharynx is largely abrogated (Avery, 1993; Darby et al., 2002; Labrousse et al., 2000), raises the possibility that this is a genuine DR mutant. Yet in terms of the indicators of reduced nutrition examined here, it is one of the least severe of the 8 mutants studied (Figure 3), arguing against this interpretation. *phm-2* longevity is also unlikely to reflect the apparent innate immune response caused by passage of live *E. coli* into the intestine (Kumar et al., 2019), since it is not suppressed by carbenicillin (Fig. 2E).

Arguably, the strongest evidence supporting the view that *eat-2* longevity is attributable to DR is that from studies of genes required for life extension. Here the literature is complex, with 10 different DR protocols showing differential dependency on a range of genes and pathways (Greer and Brunet, 2009; Kapahi et al., 2016). These include several cases where abrogation of function of a given gene suppresses longevity caused by both *eat-2* and bDR, including *pha-4* (Panowski et al., 2007), *nhr-62* (Heestand et al., 2013), and *npl-7* (Park et al., 2010).

However, a formal possibility is that loss of nicotinic acetylcholine receptor function in *eat-2* mutants activates such pathways independently of its effect on nutrition, and it is this that causes the infection independent increase in longevity. The absence of the latter in most Eat mutants, despite their evident reduced nutrition, is in line with this possibility: if reduced nutrition in *eat-2* mutants increases their lifespan, why is this not the case for many other Eat mutants with reduced nutrition? Thus, it is possible that *eat-2* mutant longevity is not attributable to actual DR, but to a DR mimetic effect, as proposed for drugs such as rapamycin and metformin, which extend lifespan by inhibiting pathways that are also inhibited by DR (Cabreiro and Gems, 2010).

### Could reduction of P death in Eat mutants be attributable to DR?

One possibility, in principle at least, is that reduced nutrition in Eat mutants enhances innate immunity, causing or contributing to reduced pharyngeal infection. Notably, in mice DR can reduce mortality from sepsis caused by bacterial infection (Hasegawa et al., 2012), and modulation of immunity has been viewed as a conserved effect of DR across taxa (Di Giosia et al., 2022). There is evidence of increased innate immunity in *phm-2(am1117)*, but this is thought to be due to live *E. coli* entering the gut lumen rather than reduced nutrition (Kumar et al., 2019). Moreover, DR by bDR (bacterial dilution) appears to extend lifespan by *suppressing* p38-regulated immunity in *C. elegans* (Wu et al., 2019). Nonetheless, suppression of P death in Eat mutants could in principle be a model system for investigating possible effects of reduced nutrition on innate immunity. However, for this purpose, *eat-6* would be a better model than *eat-2*, since the *eat-6* increase in life span is solely attributable to effects on the P subpopulation - to reduced P death frequency and increased P subpopulation lifespan (Fig. 1C, 1D right).

### Implications of *eat-2* Age not being attributable to DR

That *eat-2* Age is at least partially and perhaps wholly independent of DR has implications for the interpretation of the many studies that have used it as a DR model. First, for those in which proliferative *E. coli* was present, effects on *eat-2* Age could have involved altered P death frequency, altered p lifespan or both. Second, where effects were infection independent, there is now less certainty that *eat-2* Age is attributable to DR.

Does the fact that Eat mutants act independently of reduced nutrition to extend lifespan by protecting against bacterial infection mean that they are wholly irrelevant to DR in other organisms? Here it may be argued: not necessarily. One concern voiced about DR is that life-extension could reflect rescue from toxic effects of food rather than inhibition of the underlying aging process, at least in some cases (Walker et al., 2005). Toxic food effects include pathological effects of over-eating (e.g. via obesity) as seen in rodents (Weindruch et al., 1986), toxic constituents in fly food in *Drosophila* studies (Piper and Partridge, 2007), and bacterial pathogenicity in *C. elegans* (Garigan et al., 2002; Gems and Riddle, 2000). DR in rodents is unlikely to act by rescue from over-eating (glutton rescue) (Austad and Kristan, 2003; Nelson et al., 1985), but may well do in rhesus macaques in an experimental setting where high-fat and high-sucrose semi-purified diet was employed (Mattison et al., 2017; Mattison et al., 2012). Thus, protection against infection in *C. elegans* Eat mutants is somewhat akin to certain effects of food restriction in higher organisms insofar as they act mainly by rescue of toxic effects of food, rather than suppression of endogenous mechanisms of aging.

Another distant parallel to anti-aging effects of DR in mammals is the reduction of mechanical senescence of the pharynx in *eat-2* mutants (Chow et al., 2006; Zhao et al., 2017) and, plausibly, in Eat mutants generally (this study). However, this consequence of *eat-2* is clearly a DR effect of a very different sort to those that extend lifespan in mammals (which lack a pharyngeal cuticle).

### Condition selection bias and use of Eat mutants as DR regimen

A given experimental test can give different results, depending on how the test is designed. For example, whether inhibition of autophagy using RNAi suppresses *daf-2* insulin/IGF-1 receptor mutant longevity depends on the gene in the autophagy pathway inhibited, the *daf-2* allele used, and ambient temperature (Hsiung et al., 2026). Such experimental condition dependency creates a risk of condition selection bias, where investigators favor conditions that yield expected results (Hsiung et al., 2026). The use of *eat-2* for DR studies is a potential example of condition selection bias, that occurred in early *C. elegans* DR studies, and affected later studies in a manner that investigators were almost certainly unaware of. In other words, *eat-2* mutants were preferred because they more reliably extended lifespan, particularly in the absence of bacterial proliferation. Condition selection bias potentially contributes to difficulties with the reproducibility of experimental results, what has been called the reproducibility crisis (Prinz et al., 2011; Ritchie, 2020; Voelkl et al., 2020).

### The challenge of distinguishing reduced nutrition from DR

This study draws attention to a somewhat complex problem of interpretation, relating to the distinction between reduced nutrition and DR. Our finding that many Eat mutants show clear indications of reduced nutrition but are not long-lived presents a challenge for future *C. elegans* studies that attempt to attribute longevity to a DR effect. This is because many markers of reduced nutrition (e.g. mRNA profile signatures or *acs-2::GFP* expression (Burkewitz et al., 2015)) will inevitably be detectable in both the presence and absence of life-extension. Thus, use of markers of reduced nutrition will need to interpreted with caution when attempting, for example, to establish whether or not infection-independent *eat-2* longevity is attributable to DR. Similarly, if it were demonstrated that a pathway mediating DR effects, such as the mTOR (mechanistic target of rapamycin) pathway (Kennedy and Lamming, 2016), is altered in *eat-2* mutants, it would remain unclear whether such an alteration is or is not mediated by reduced nutrition.

A further question raised by this study is why, in most cases, does reduced nutrition in Eat mutants not extend lifespan? After all, reduced nutrition resulting e.g. from bacterial dilution (bDR) is sufficient to increase lifespan, even where bacterial proliferation is blocked using antibiotics (Wu et al., 2019). One difference between Eat mutants and bDR is that in the former nutrition is reduced throughout development as well as during adulthood, whereas in the latter it is reduced during adulthood alone. This could imply that to see a life-extending DR effect in *C. elegans*, food restriction needs to be limited to adulthood.

## Conclusions

Given infection-dependent effects at least, re-consideration is warranted for conclusions drawn from previous studies that have employed *eat-2* mutants as a model for DR, including many of those listed in Supplementary Table 1. *eat-2* mutants are most often used in epistasis-type experiments, to test whether a given treatment (e.g. mutation, drug) involves a mechanism shared with that by which DR extends lifespan. Any observed interaction could involve either infection-dependent mechanisms that affect P death frequency, or P death- independent mechanisms that affect p Age, or a combination of the two. Careful reinterpretation of prior findings with *eat-2*, supported by further investigation, has the potential to draw new and clearer insights from a substantial body of previous work on *C. elegans* aging.

To emphasize again: the original reason for believing that *eat-2* Age is attributable to DR is that Eat mutants are typically long-lived (Lakowski and Hekimi, 1998). This study shows that Eat mutant longevity, though typical of this mutant class, is usually not attributable to DR. While this does not exclude the possibility that infection-independent *eat-2* Age is attributable to DR, it removes the original reason for believing this in the first place. However, the widespread adoption of *eat-2* as a DR model raises the burden of proof in this case. Thus, further investigation is warranted to establish the cause of *eat-2* Age. In short, although our study does not exclude the possibility that *eat-2* mutant longevity reflects DR, it does raise doubts about it.

## Experimental Methods

### Culture conditions and strains

*C. elegans* maintenance was performed using standard protocols (Brenner, 1974). Strains were grown at 20°C on nematode growth medium (NGM) agar plates seeded with *E. coli* OP50. *C. elegans* strains used included: N2 (hermaphrodite stock) (Zhao et al., 2019), DA531 *eat-1(ad427) IV; fln-2(ot611) X*, GA6005 *eat-1(ad427)*, DA465 *eat-2(ad465)*, DA1113 *eat-2(ad1113)*, DA1116 *eat-2(ad1116)*, DA464 *eat-5(ad464)*, DA467 *eat-6(ad467)*, DA606 *eat-10(ad606)*, DA1110 *eat-18(ad1110)*, DA597 *phm-2(ad597)*, DA493 *phm-3(ad493) III; fln-2(ot611) X*, GA6003 *phm-3(ad493)*.

### Lifespan assays

Unless otherwise specified, lifespan assays were performed at 20°C on NGM plates with a 2-day old OP50 lawn. FUDR (5-fluorodeoxyuridine) was not used in this study. Worms were transferred daily during the reproductive period to remove progeny, and every 3-7 days thereafter. Worms lost due to causes other than aging, e.g. internal hatching (bagging) or desiccation on the plate wall, were right censored. Mortality and pharyngeal morphology of corpses were scored every 2-3 days. For antibiotic treatment, 100 μl of 500 mM carbenicillin was added to 2-day old lawns of OP50 and allowed to dry overnight before use (final concentration 4 mM). Survival plots were generated in GraphPad Prism using combined lifespan data, with L4 as day 0 on the time scale.

### Microscopy

For the pharyngeal infection assay, worms were fed *E. coli* OP50 expressing red fluorescent protein (OP50-RFP), and allowed to crawl on non-fluorescent OP50 plates for 5 minutes to remove excess fluorescent bacteria before examination. Live worms were anesthetized in 0.2% levamisole, and mounted on 2% agarose pads. For Nomarski and epifluorescence microscopy, slides were imaged using a Zeiss ApoTome.2 microscope system, fitted with a Hamamatsu C13440 ORCA-Flash4.0 V3 digital camera. Zen software was used for image acquisition.

### Brood size measurement

Individual L4 larvae were placed on NGM plates seeded with OP50, and worms were transferred every 24 hr until reproduction ceased, and the number of progeny were recorded each day. A minimum of 10 brood per strain were scored.

### Lawn avoidance measurement

NGM plates with small, circular lawns were prepared by pipetting 100 μl overnight culture of OP50 onto the plate center and allowed to dry. NGM plates with large lawns, covering the plate surface, were prepared similarly but using 1 ml of OP50 culture and rocking the plate to ensure coverage of the entire surface. For carbenicillin-treated lawns, 100 μl of 500 mM carbenicillin was added around the lawn 24 hr after seeding, and allowed to dry overnight. 50 - 100 eggs were transferred onto the bacteria lawn taking care to avoid disrupting it, and incubated at 20°C. Nematodes were then scored and transferred every 24 hr over a 4 day period. Avoidance was calculated as the percentage of animals outside of the lawn, i.e. the number outside the lawn divided by the total number of animals.

### Statistics

Survival analyses were conducted using GraphPad Prism, and differences in survival were tested using the non-parametric log rank test. Raw mortality data are available in Supplemental Dataset 1. An unpaired Student’s *t*-test was used to compare frequencies of P death between different strains and/or conditions. Linear regression analyses were performed using GraphPad Prism to test whether slopes are significantly different to zero. The Wilcoxon signed rank test was used to compare infection frequencies between pharygeal quadrants. No statistical methods were used to predetermine sample size. The experiments were not randomized. The investigators were not blinded to allocation during experiments and outcome assessment.

## Acknowledgments

We thank A.V. Ha, C. Macleod, G. Murphy, E. Strachan, S. Taylor and S. Wang for technical assistance. We would like to thank F. Cabreiro, J. Labbadia and members of the Gems laboratory for useful discussion. Some strains were provided by the Caenorhabditis Genetics Center, which is funded by NIH Office of Research Infrastructure Programs (P40 OD010440).

## Supplementary Information

Contents Summary

## Supplementary figures

**Supplementary Figure 1.**
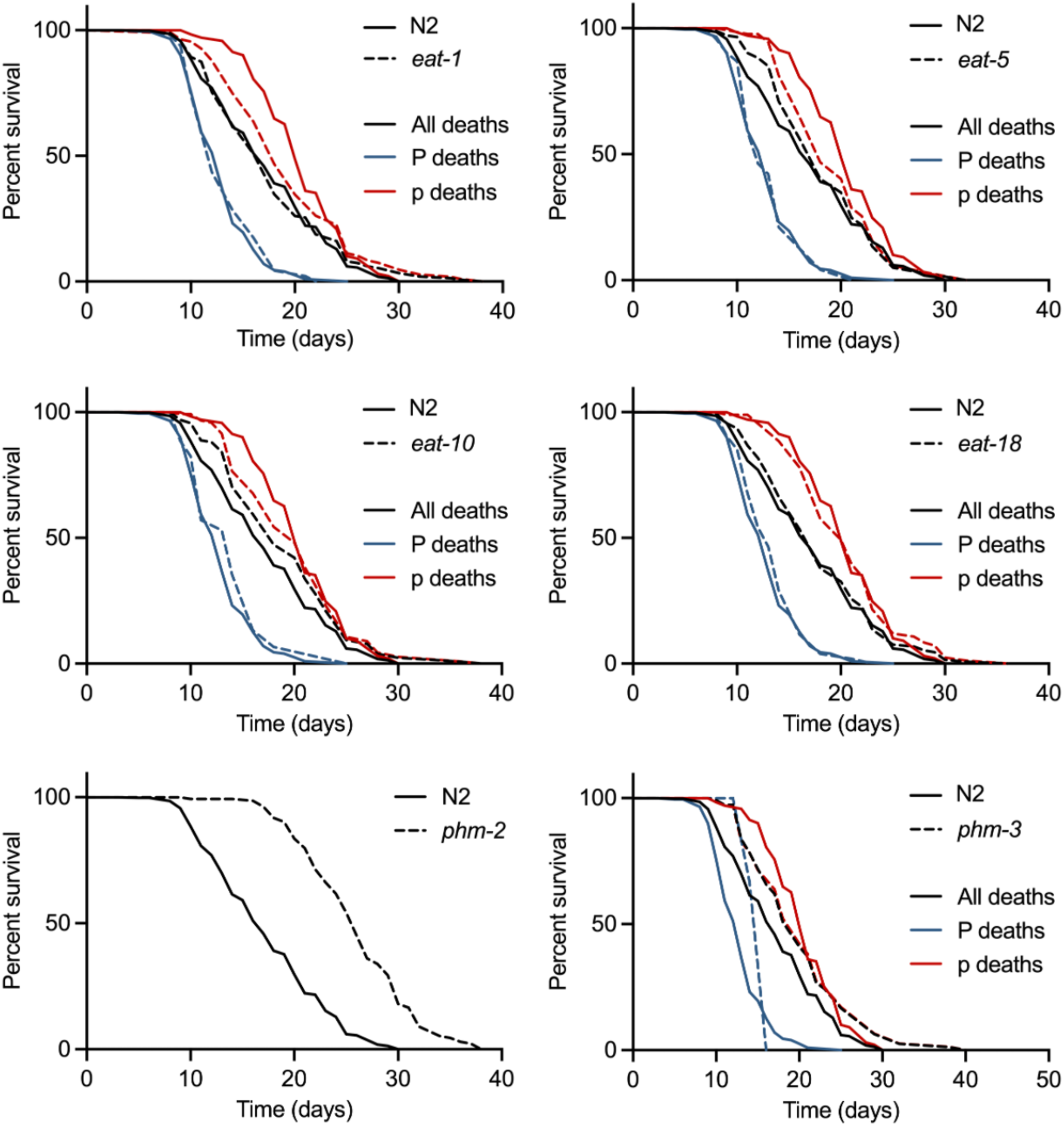
Mortality deconvolution data for selected Eat mutants. Survival curves of the whole population, the P sub-population and the p sub-population.

**Supplementary Figure 2.**
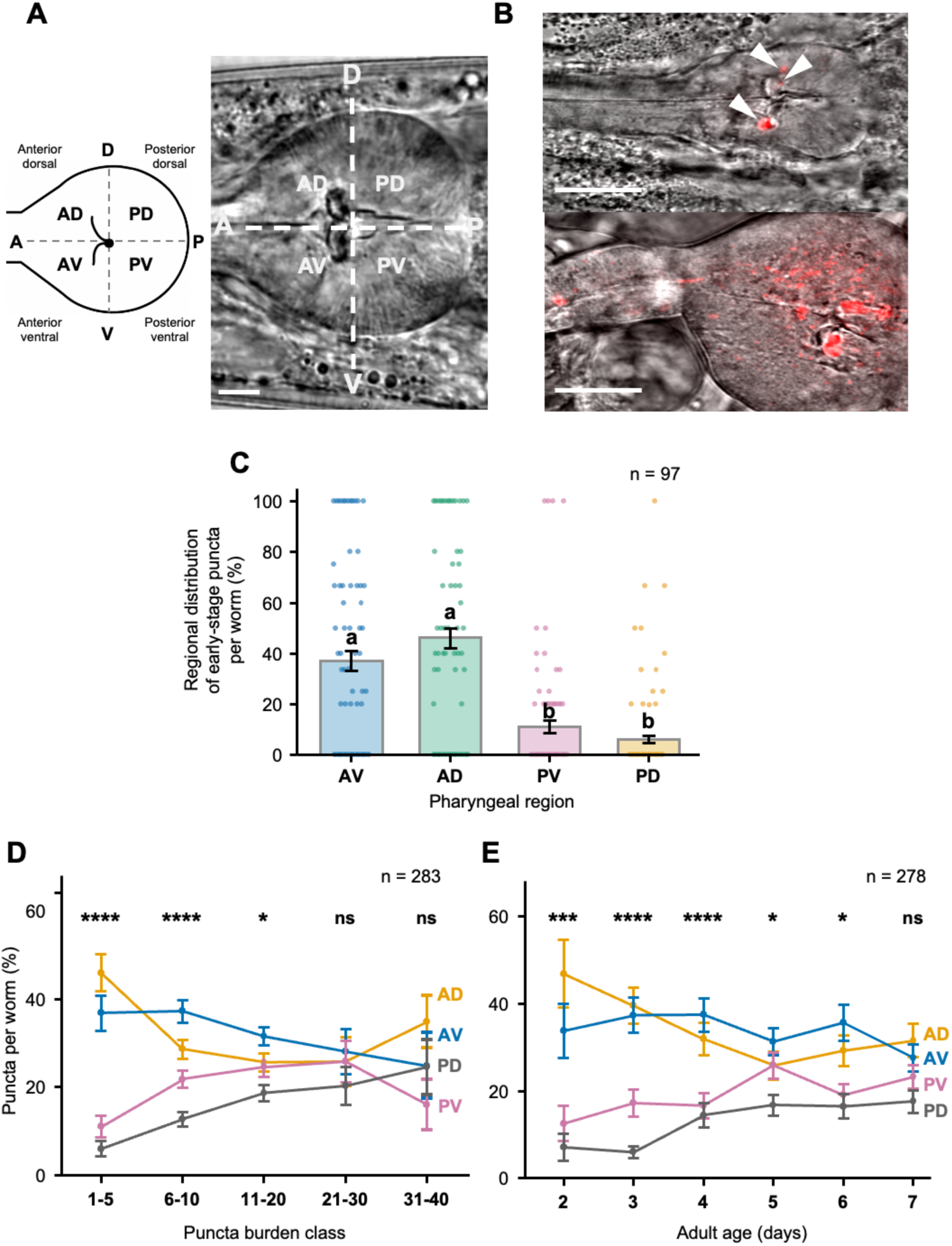
Site of initiation of infection in the pharyngeal bulb anterior. (**A**) Four quadrants of the terminal pharyngeal bulb: AV, anterior ventral, AD, anterior dorsal, PV, posterior ventral, PD, posterior dorsal. Left, schematic representation of the pharyngeal bulb. Right, brightfield image (Nomarski). Scale bar: 10 μM. (**B**) Example of pharyngeal infection with *E. coli* OP50 expressing RFP. Top, early stage infection (day 5 of adulthood). Arrowheads: localized infection visible as red fluorescent puncta in anterior dorsal and anterior ventral quadrants (2 and 1 puncta/punctum, respectively). Bottom, example of advanced stage infection (day 9 of adulthood, live individual), where numerous individual puncta have coalesced, such that counting them is no longer practicable. Note the swollen state of the pharynx, preceding P death. Merged bright field and epifluorescence images. Scale bar: 20 μM. (**C**) Early-stage infection (1–5 discrete RFP puncta) in the four pharyngeal quadrants, showing elevated frequency in anterior quadrants (*n* = 97 animals, mainly D2-D4 of adulthood). (**D**) Puncta number differs between pharyngeal quadrants only at lower puncta frequencies (mostly D1-D7 animals). (**E**) Puncta number differs between pharyngeal quadrants only at earlier ages. Individuals with swollen, pre-P death pharynxes were not included in this analysis, due to puncta no longer being countable. (**C**-**E**) ns *p* > 0.05, * *p* ≤ 0.05, ** *p* ≤ 0.01, *** *p* ≤ 0.001, **** *p* ≤ 0.0001, pairwise comparisons using Holm-corrected Wilcoxon signed-rank test.

**Supplementary Figure 3.**
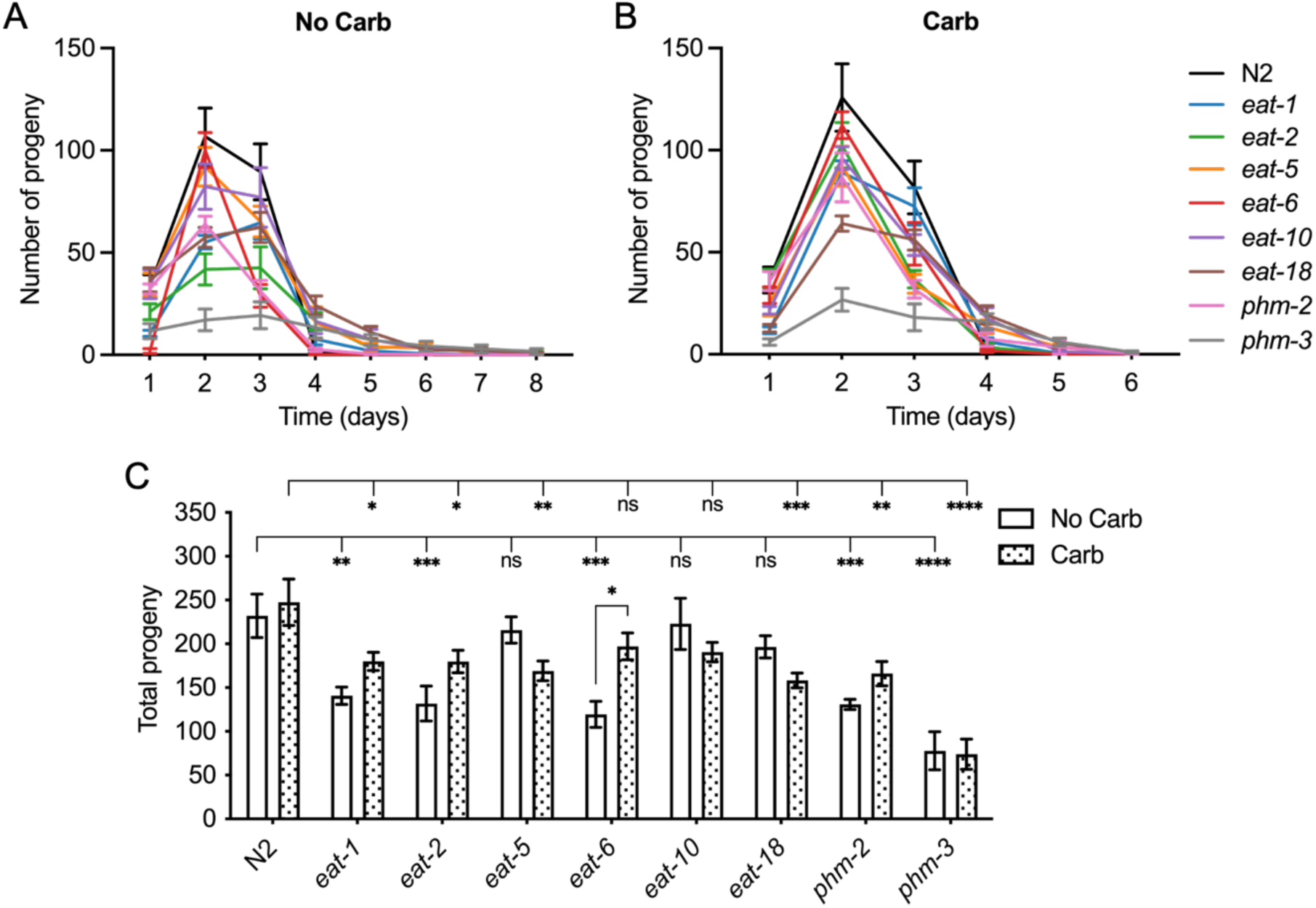
**Carbenicillin has little effect on reduced reproduction in Eat mutants**. (**A, B**) Reproductive schedule, no Carb (**A**), Carb (**B**). (**C**) Brood size. Dunnett’s multiple comparisons test, \**p*<0.05, \*\**p*<0.01, \*\*\**p*<0.001, \*\*\*\**p*<0.0001. n=7-10 for each strain.

**Supplementary Figure 4.**
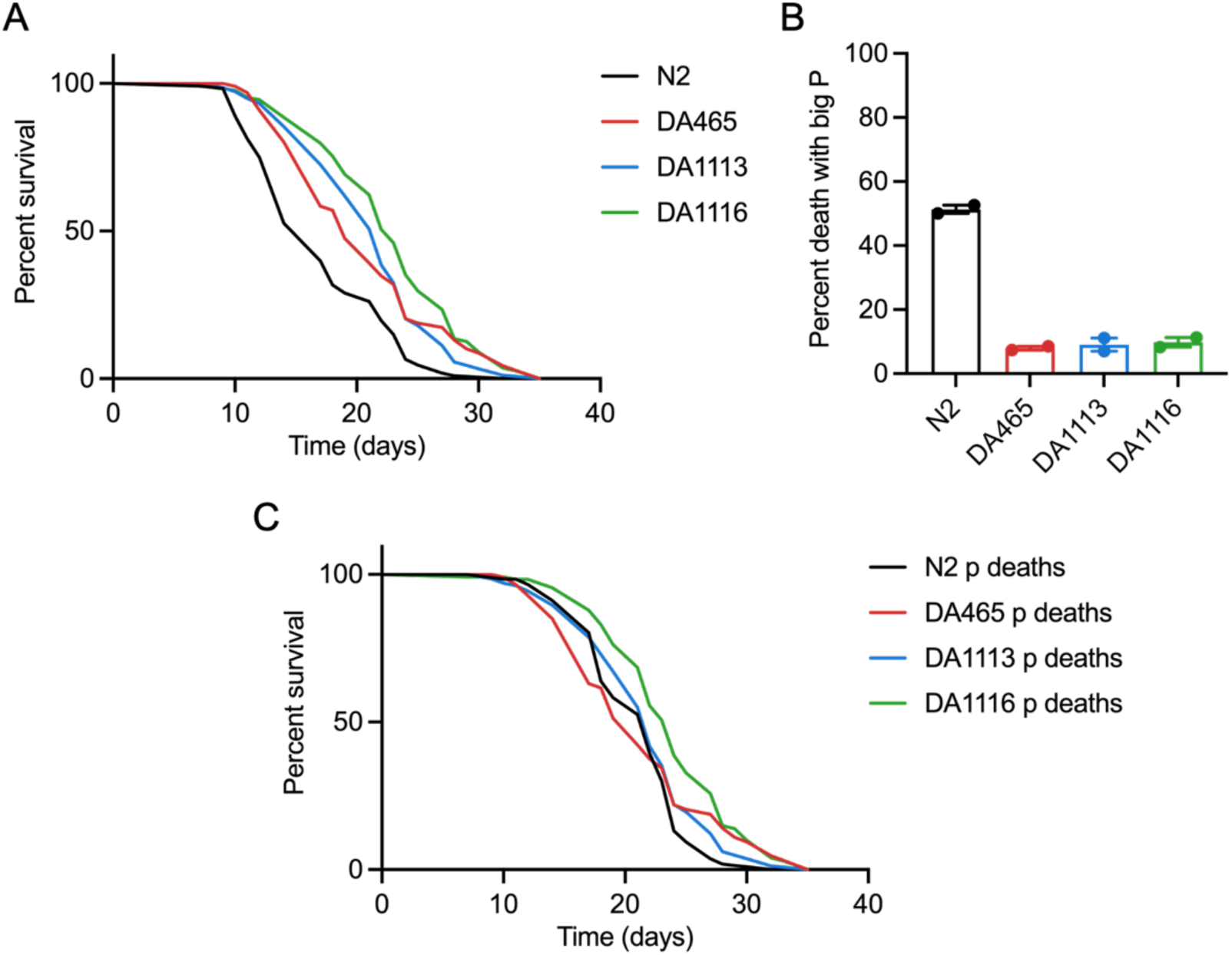
Effects of 3 *eat-2* alleles on overall and p subpopulation lifespan, and P death frequency. Culture on proliferating *E. coli* OP50. (**A**) Lifespans of overall populations, pooled data. (**B**) P death frequency, pooled data. (**C**) Lifespans of p subpopulations, pooled data. *N* = 2.

**Supplementary Figure 5.**
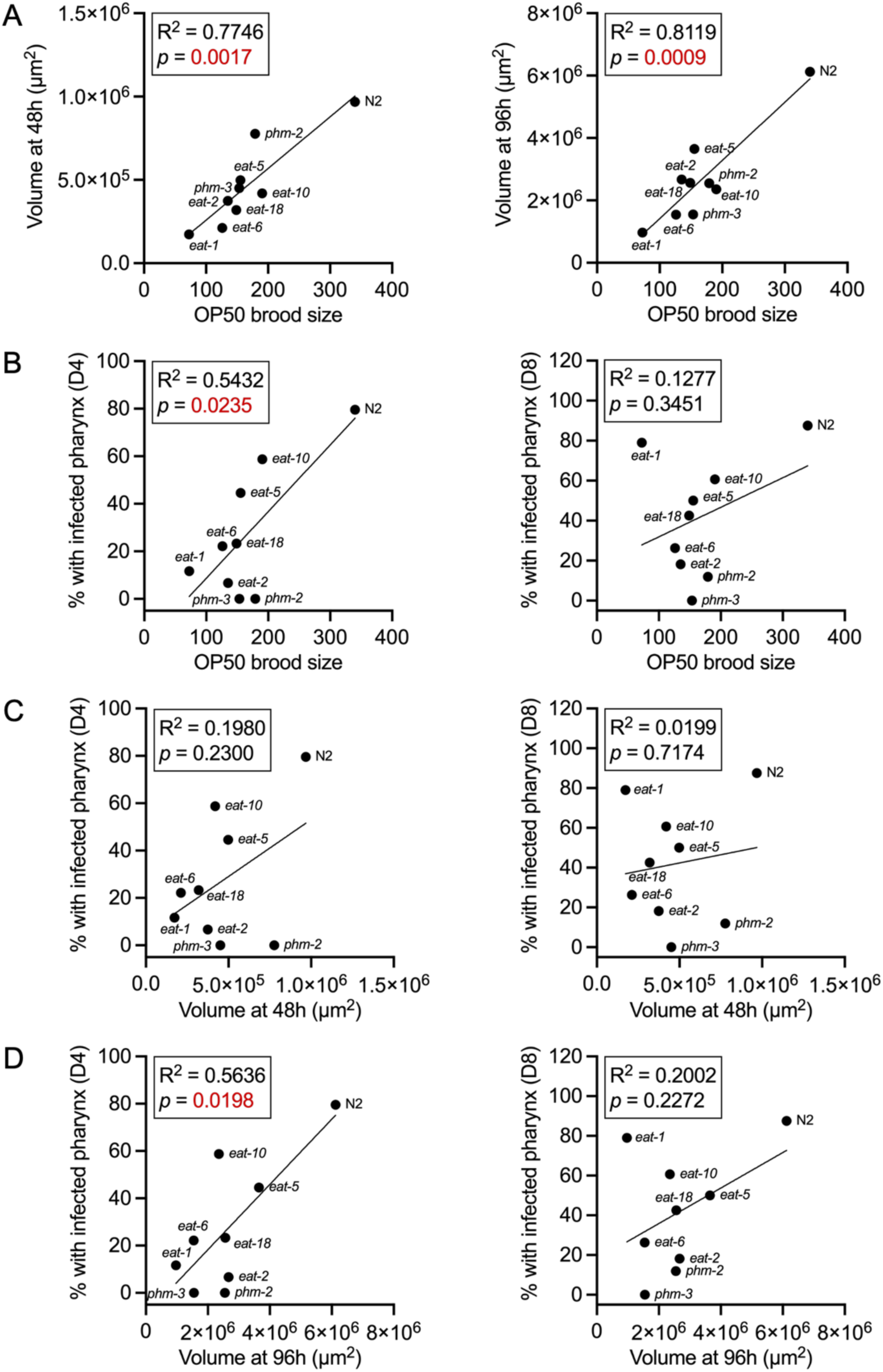
Correlation between metrics of malnutrition and early bacterial invasion in Eat mutants. (**A**) Correlations between body size (estimated volume) and brood size. (**B**) Correlation between brood size and either early bacterial infection on day 4 (left) or day 8 (right). (**C**) Correlation between 48h body size and either early bacterial invasion on day 4 (left) or day 8 (right). (**D**) Correlation between 96h body size and either early bacterial invasion on day 4 (left) or day 8 (right).

**Supplementary Figure 6.**
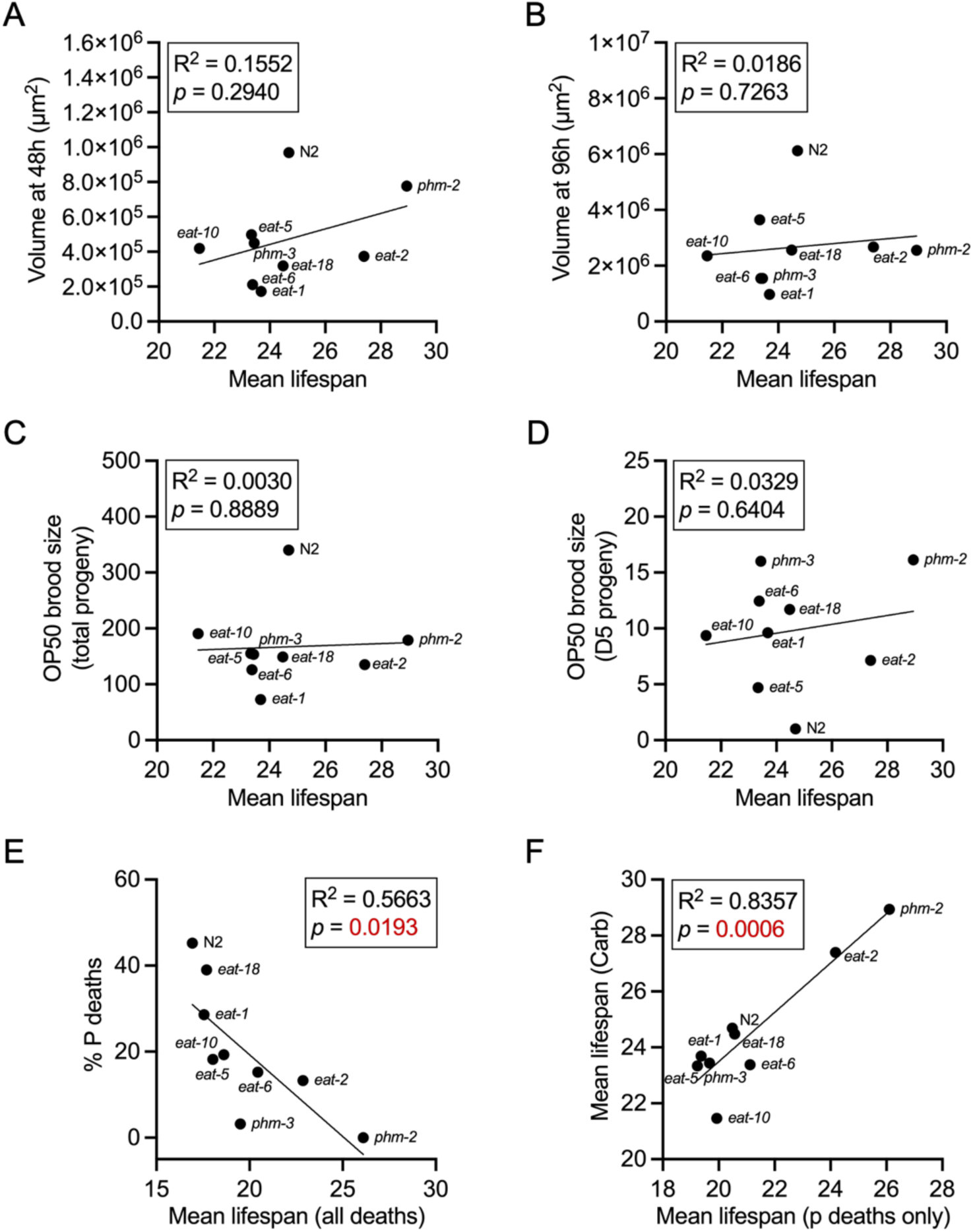
Correlation between mean lifespan in carbenicillin-treated worms with metrics of malnutrition. (**A**) Correlation with body size (estimated volume) at 48h after egg lay. (**B**) Correlation with body size (estimated volume) at 96h after egg lay. (**C**) Correlation with brood size. (**D**) Correlation with Day 5 progeny count. (**E**) Correlation between overall lifespan with % P deaths. (**F**) Correlation of lifespan between p populations (no carbenicillin) and carbenicillin-treated populations.

**Supplementary Figure 7.**
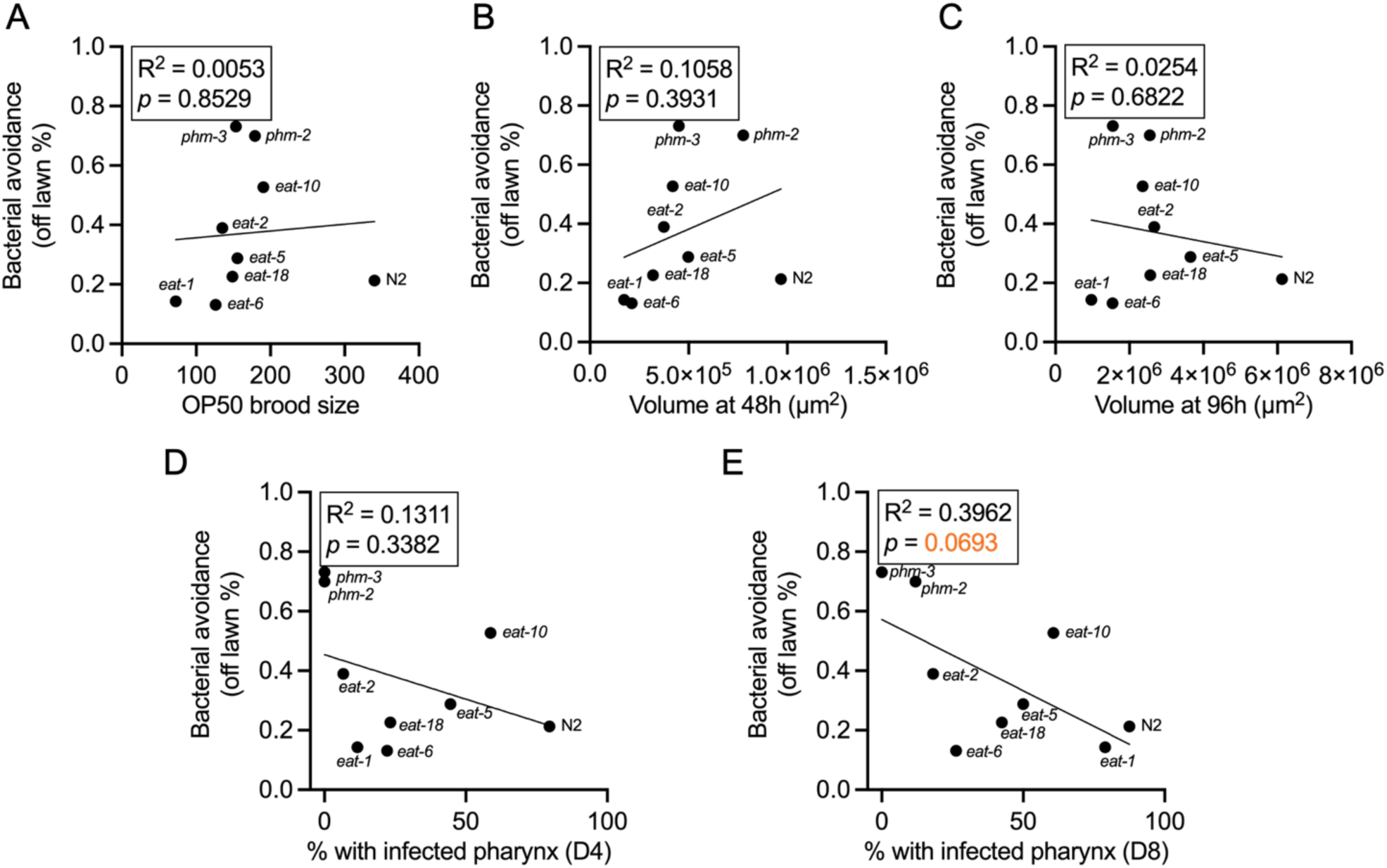
Regression analysis of lawn avoidance. (**A**) Correlation with brood size. (**B**) Correlation with body size (estimated volume) at 48h after egg lay. (**C**) Correlation with body size (estimated volume) at 96h after egg lay. (**D, E**) Correlation with bacterial invasion on day 4 (**D**) and day 8 (**E**).

**Supplementary Figure 8.**
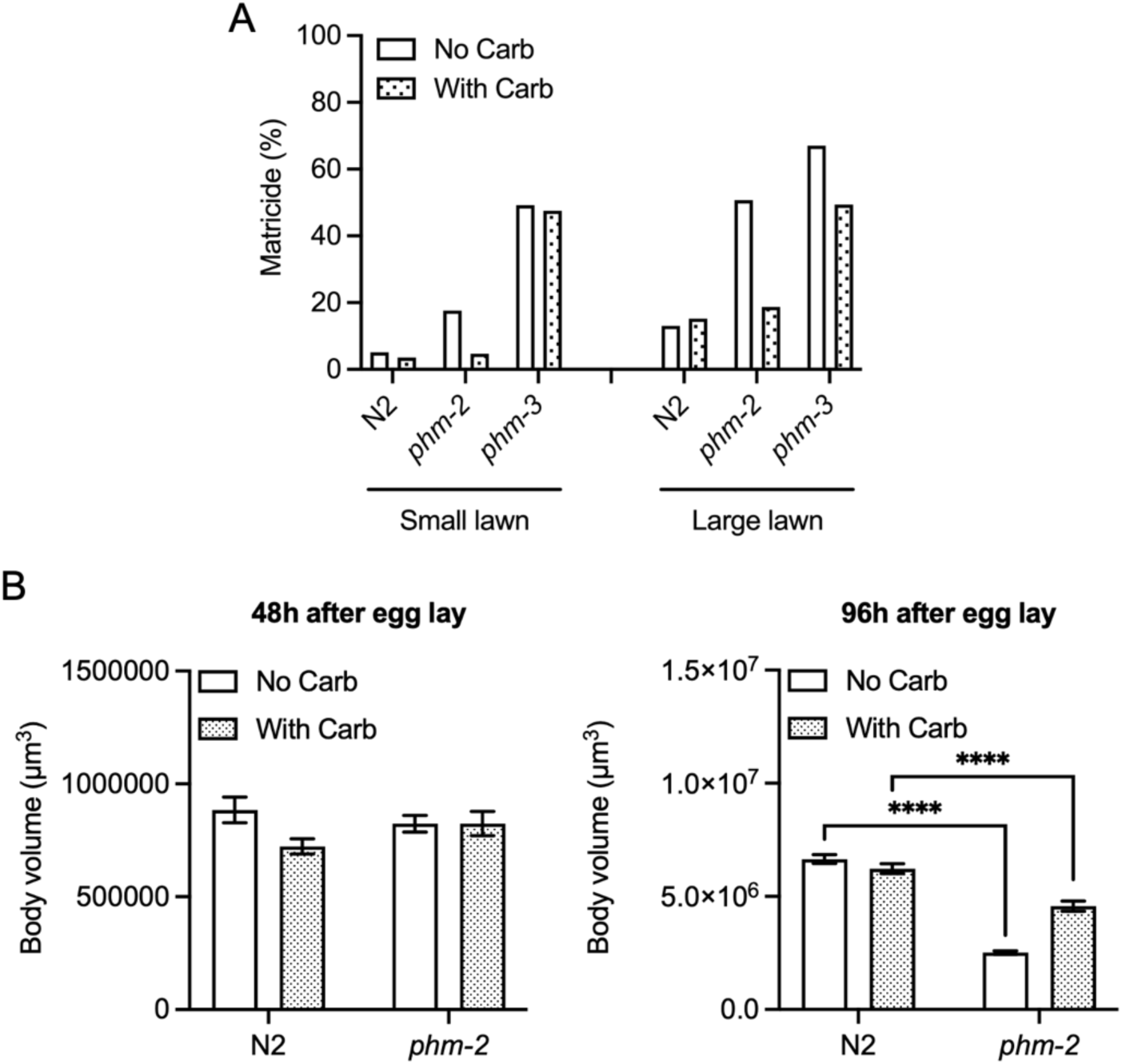
Effects of carbenicillin on *phm-2(ad597)* mutants on large lawns. (**A**) Carbenicillin suppresses elevated levels of matricide in *phm-2* mutants on large lawns. (**B**) Carbenicillin rescues the reduction in body size at 96 hr in *phm-2* mutants on large lawns. Thus, on large lawns proliferative *E. coli* causes matricide and reduces growth in *phm-2* mutants.

**Supplementary Table 1.**
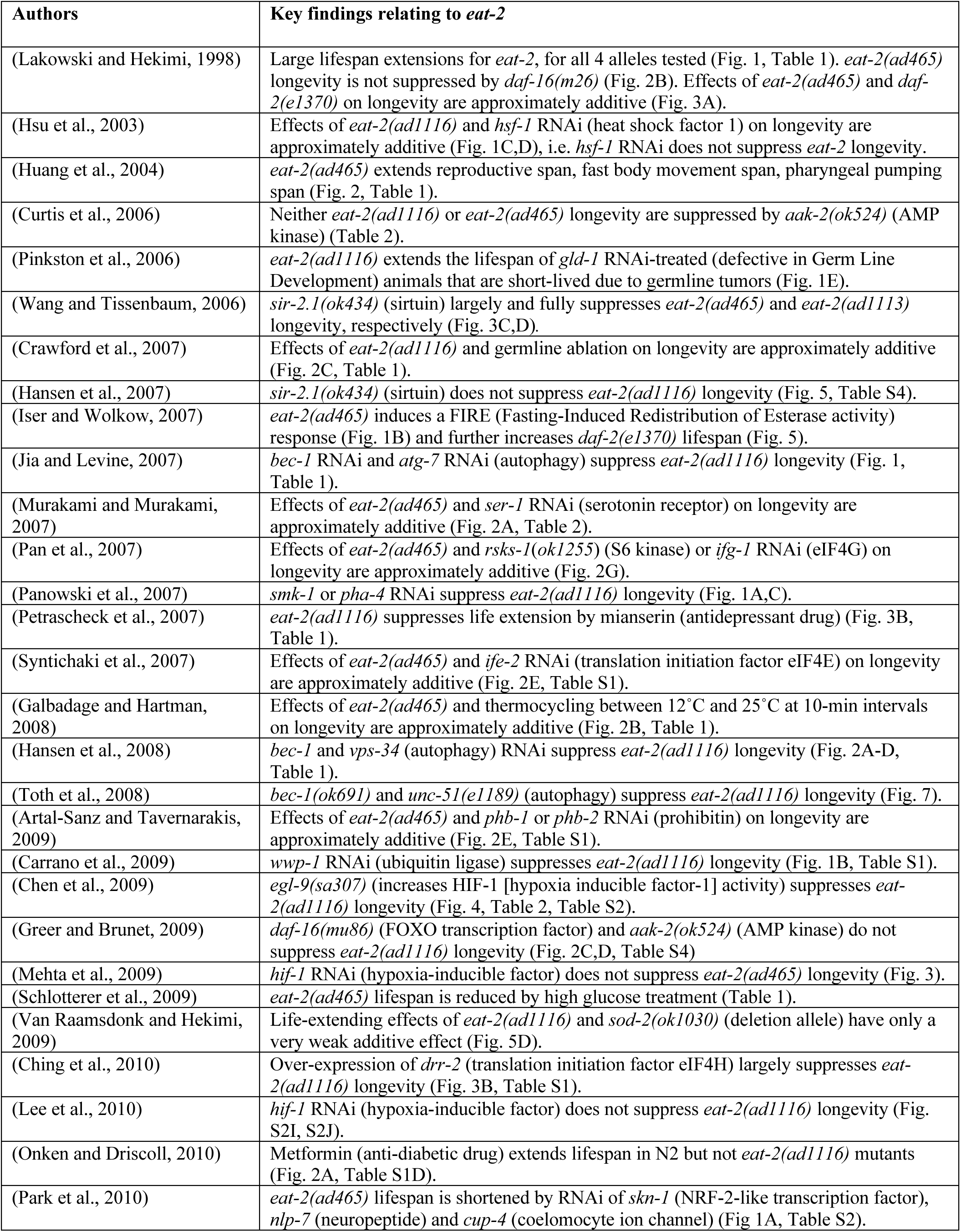
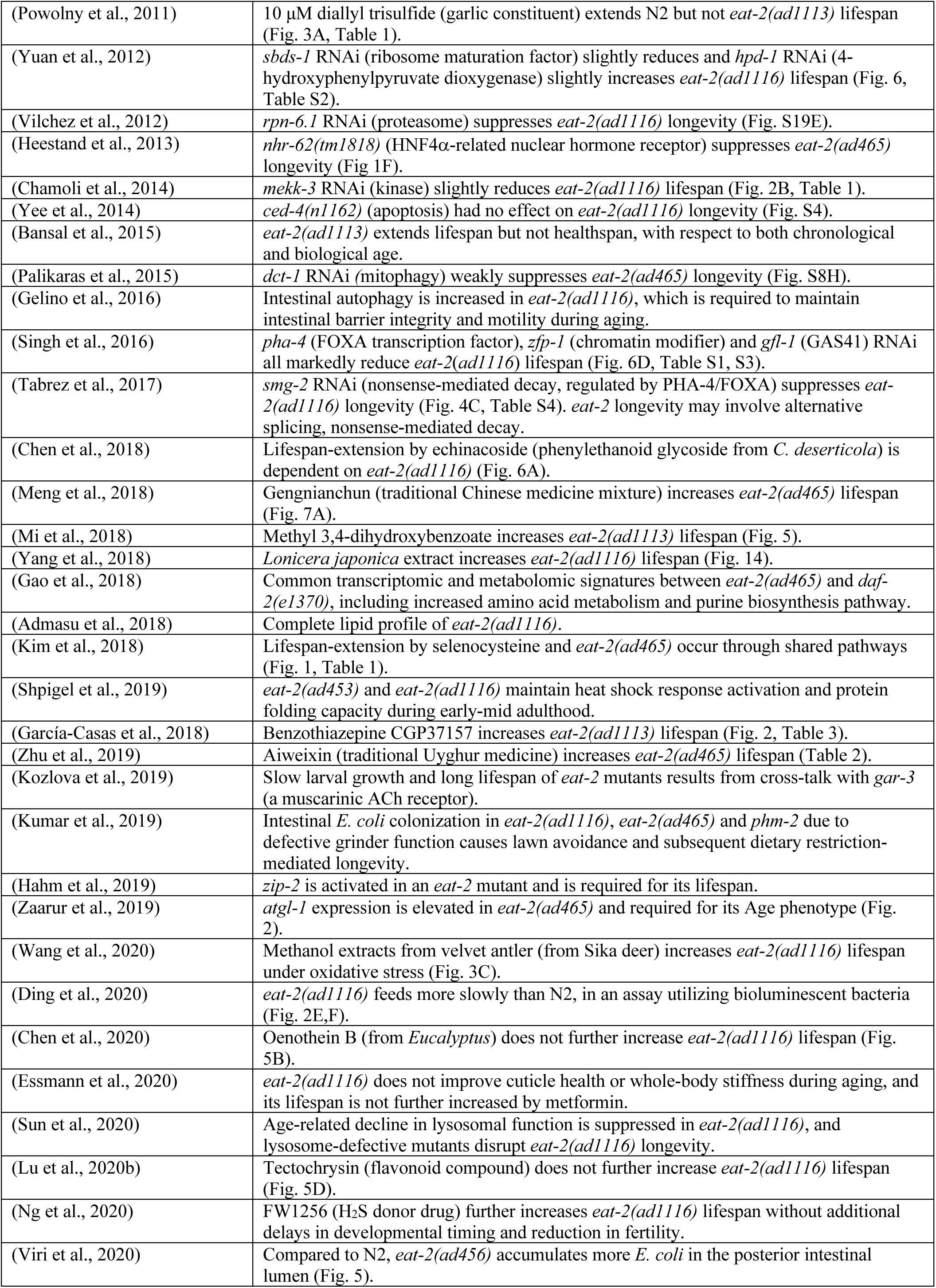
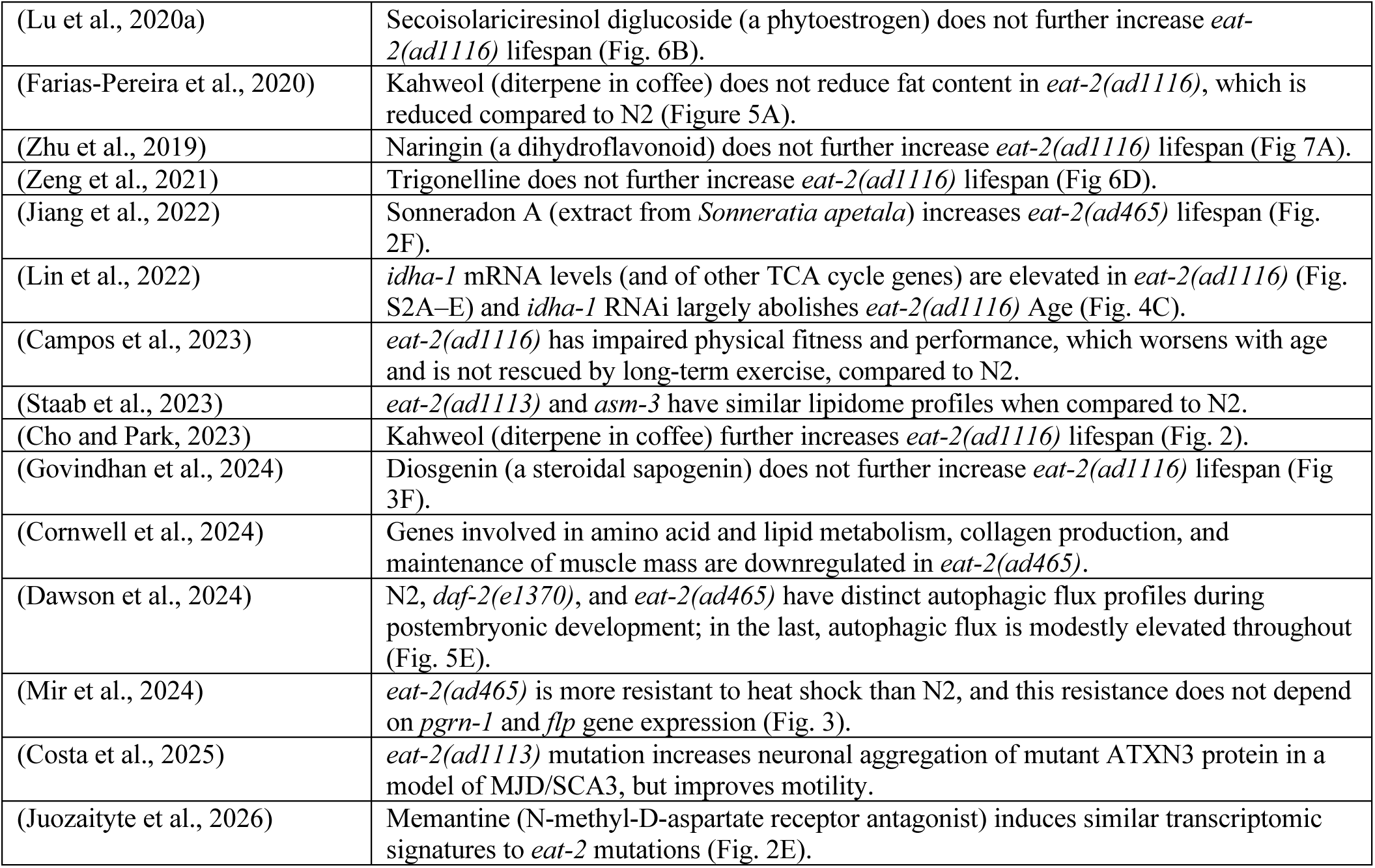
Findings from 77 previous studies employing *eat-2* mutants as a model for dietary restriction.

**Supplementary Table 2.**
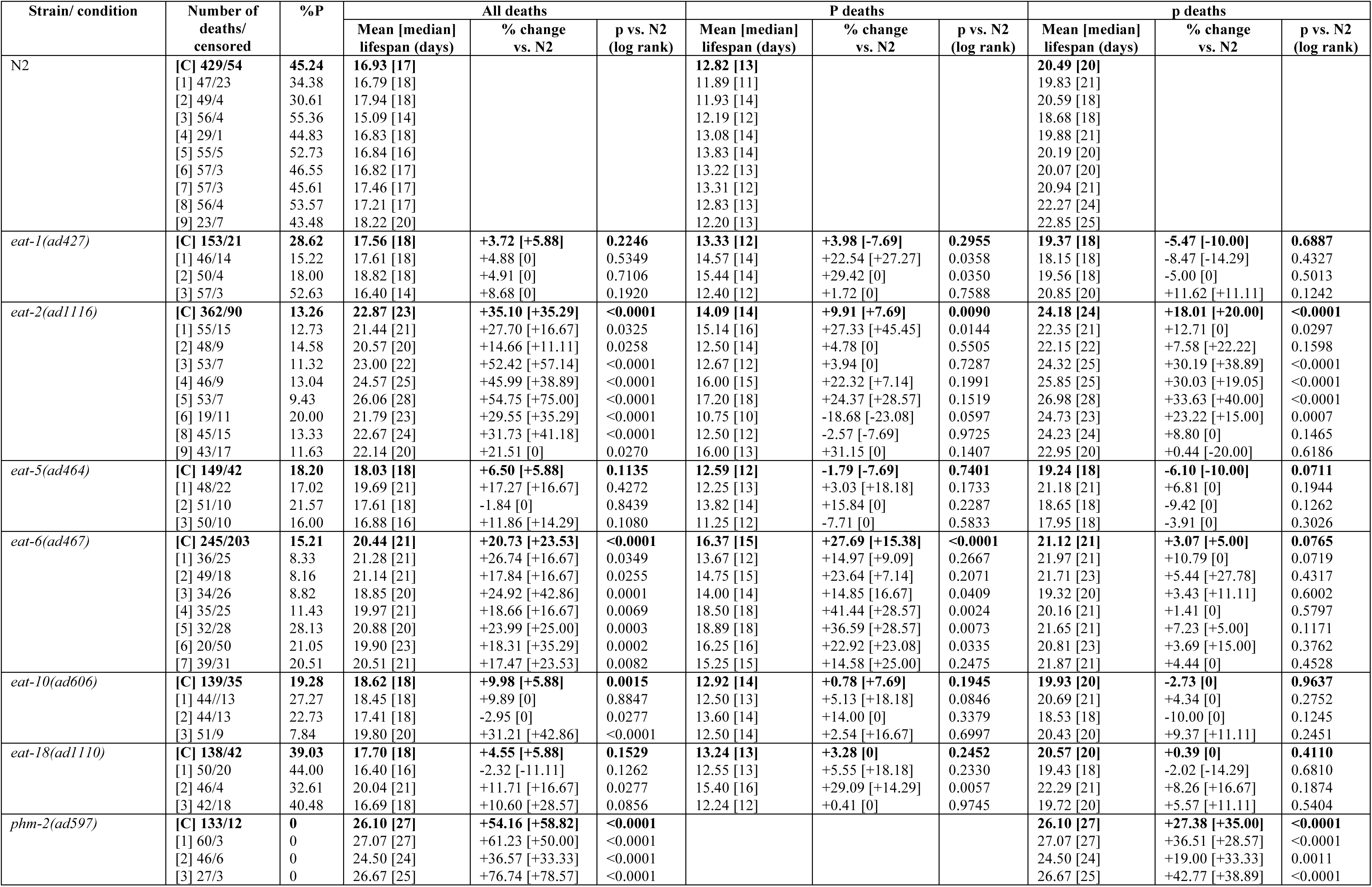
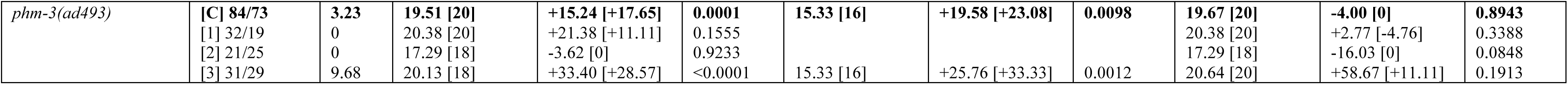
Lifespan analysis of Eat mutants on proliferating bacteria (20°C)

**Supplementary Table 3.** Lifespan analysis of Eat mutants on non-proliferating bacteria (20°C)

| Strain/ condition | Number of deaths/censored | Mean [median] lifespan (days) | % change vs. N2 | p vs. N2 (log rank) |
| --- | --- | --- | --- | --- |
| N2 | [C] 199/9<br>[1] 44/1<br>[2] 53/4<br>[3] 45/1<br>[4] 57/3 | 24.69 [24]<br>23.80 [24]<br>23.26 [23]<br>24.84 [26]<br>26.58 [26] |  |  |
| *eat-1(ad427)* | [C] 137/12<br>[1] 51/1<br>[3] 29/8<br>[4] 57/3 | 23.69 [24]<br>23.75 [23]<br>25.69 [26]<br>22.61 [24] | -4.05 [0]<br>-0.21 [-4.17]<br>+3.42 [0]<br>-14.94 [-7.69] | 0.0518<br>0.7648<br>0.6275<br>0.0029 |
| *eat-2(ad1116)* | [C] 192/15<br>[1] 45/2<br>[2] 25/3<br>[3] 63/9<br>[4] 59/1 | 27.40 [28]<br>26.62 [26]<br>26.68 [26]<br>28.22 [28]<br>27.42 [28] | +10.98 [+16.67]<br>+11.85 [+8.33]<br>+14.70 [+13.04]<br>+13.61 [+7.69]<br>+3.16 [+7.69] | 0.0078<br>0.0620<br>0.1815<br>0.0115<br>0.8892 |
| *eat-5(ad464)* | [C] 190/31<br>[1] 49/4<br>[2] 52/3<br>[3] 44/9<br>[4] 45/15 | 23.34 [23]<br>21.04 [18]<br>19.90 [19]<br>27.14 [28]<br>26.11 [26] | -5.47 [-4.17]<br>-11.60 [-25.00]<br>-14.45 [-17.39]<br>+9.26 [+7.69]<br>-1.77 [0] | 0.0816<br>0.1128<br>0.0050<br>0.1321<br>0.7414 |
| *eat-6(ad467)* | [C] 196/40<br>[1] 47/1<br>[2] 57/4<br>[3] 57/10<br>[4] 35/25 | 23.38 [23]<br>21.00 [18]<br>22.44 [21]<br>26.70 [26]<br>22.71 [21] | -5.31 [-4.17]<br>-11.76 [-25.00]<br>-3.53 [-8.70]<br>+7.49 [0]<br>-14.56 [-19.23] | 0.0221<br>0.0958<br>0.6250<br>0.3194<br>0.0020 |
| *eat-10(ad606)* | [C] 162/40<br>[1] 56/1<br>[2] 46/3<br>[3] 28/8<br>[4] 32/28 | 21.46 [22]<br>20.63 [22]<br>20.20 [19]<br>24.54 [26]<br>22.06 [21] | -13.08 [-8.33]<br>-13.32 [-8.33]<br>-13.16 [-17.39]<br>-1.21 [0]<br>-17.01 [-19.23] | <0.0001<br>0.1648<br>0.0135<br>0.5546<br>0.0082 |
| *eat-18(ad1110)* | [C] 203/39<br>[1] 32/1<br>[2] 47/5<br>[3] 82/15<br>[4] 42/18 | 24.48 [24]<br>25.41 [25]<br>19.77 [19]<br>27.40 [28]<br>23.36 [24] | -0.85 [0]<br>+6.76 [+4.17]<br>-15.00 [-17.39]<br>+10.30 [+7.69]<br>-12.11 [-7.69] | 0.5378<br>0.6891<br>0.0019<br>0.0441<br>0.0065 |
| *phm-2(ad597)* | [C] 227/11<br>[1] 54/1<br>[2] 47/3<br>[3] 70/3<br>[4] 56/4 | 28.94 [28]<br>29.83 [30]<br>28.53 [30]<br>30.29 [30]<br>26.75 [28] | +17.21 [+16.67]<br>+25.34 [+25.00]<br>+22.66 [+30.43]<br>+21.94 [+15.38]<br>+0.64 [+7.69] | <0.0001<br><0.0001<br>0.0056<br><0.0001<br>0.5769 |
| *phm-3(ad493)* | [C] 174/53<br>[1] 43/1<br>[2] 51/1<br>[3] 54/17<br>[4] 26/34 | 23.44 [23]<br>22.67 [23]<br>22.10 [21]<br>23.65 [23]<br>26.88 [26] | -5.06 [-4.17]<br>-4.75 [-4.17]<br>-4.99 [-8.70]<br>-4.79 [-11.54]<br>+1.13 [0] | 0.0247<br>0.0694<br>0.2067<br>0.2447<br>0.7480 |

**Supplementary Table 4.**
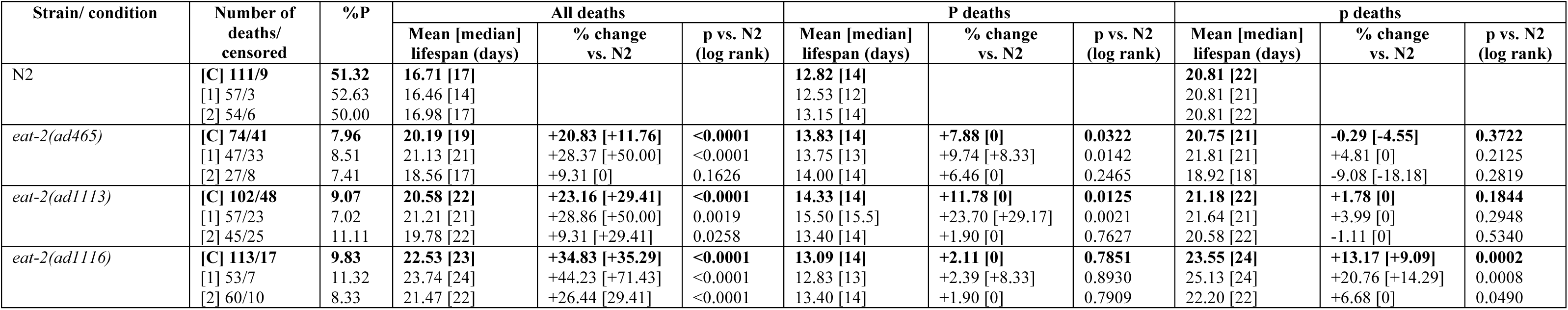
Lifespan analysis of 3 *eat-2* mutants on proliferating bacteria (20°C)

| Strain/ condition | Number of deaths/<br>censored | %P | All deaths |  |  | P deaths |  |  | p deaths |  |  |
| --- | --- | --- | --- | --- | --- | --- | --- | --- | --- | --- | --- |
|  |  |  | Mean [median]<br>lifespan (days) | % change<br>vs. N2 | p vs. N2<br>(log rank) | Mean [median]<br>lifespan (days) | % change<br>vs. N2 | p vs. N2<br>(log rank) | Mean [median]<br>lifespan (days) | % change<br>vs. N2 | p vs. N2<br>(log rank) |
| N2 | [C] 111/9<br>[1] 57/3<br>[2] 54/6 | 51.32<br>52.63<br>50.00 | 16.71 [17]<br>16.46 [14]<br>16.98 [17] |  |  | 12.82 [14]<br>12.53 [12]<br>13.15 [14] |  |  | 20.81 [22]<br>20.81 [21]<br>20.81 [22] |  |  |
| *eat-2(ad465)* | [C] 74/41<br>[1] 47/33<br>[2] 27/8 | 7.96<br>8.51<br>7.41 | 20.19 [19]<br>21.13 [21]<br>18.56 [17] | +20.83 [+11.76]<br>+28.37 [+50.00]<br>+9.31 [0] | <0.0001<br><0.0001<br>0.1626 | 13.83 [14]<br>13.75 [13]<br>14.00 [14] | +7.88 [0]<br>+9.74 [+8.33]<br>+6.46 [0] | 0.0322<br>0.0142<br>0.2465 | 20.75 [21]<br>21.81 [21]<br>18.92 [18] | -0.29 [-4.55]<br>+4.81 [0]<br>-9.08 [-18.18] | 0.3722<br>0.2125<br>0.2819 |
| *eat-2(ad1113)* | [C] 102/48<br>[1] 57/23<br>[2] 45/25 | 9.07<br>7.02<br>11.11 | 20.58 [22]<br>21.21 [21]<br>19.78 [22] | +23.16 [+29.41]<br>+28.86 [+50.00]<br>+9.31 [+29.41] | <0.0001<br>0.0019<br>0.0258 | 14.33 [14]<br>15.50 [15.5]<br>13.40 [14] | +11.78 [0]<br>+23.70 [+29.17]<br>+1.90 [0] | 0.0125<br>0.0021<br>0.7627 | 21.18 [22]<br>21.64 [21]<br>20.58 [22] | +1.78 [0]<br>+3.99 [0]<br>-1.11 [0] | 0.1844<br>0.2948<br>0.5340 |
| *eat-2(ad1116)* | [C] 113/17<br>[1] 53/7<br>[2] 60/10 | 9.83<br>11.32<br>8.33 | 22.53 [23]<br>23.74 [24]<br>21.47 [22] | +34.83 [+35.29]<br>+44.23 [+71.43]<br>+26.44 [29.41] | <0.0001<br><0.0001<br><0.0001 | 13.09 [14]<br>12.83 [13]<br>13.40 [14] | +2.11 [0]<br>+2.39 [+8.33]<br>+1.90 [0] | 0.7851<br>0.8930<br>0.7909 | 23.55 [24]<br>25.13 [24]<br>22.20 [22] | +13.17 [+9.09]<br>+20.76 [+14.29]<br>+6.68 [0] | 0.0002<br>0.0008<br>0.0490 |

**Supplementary Table 5.**
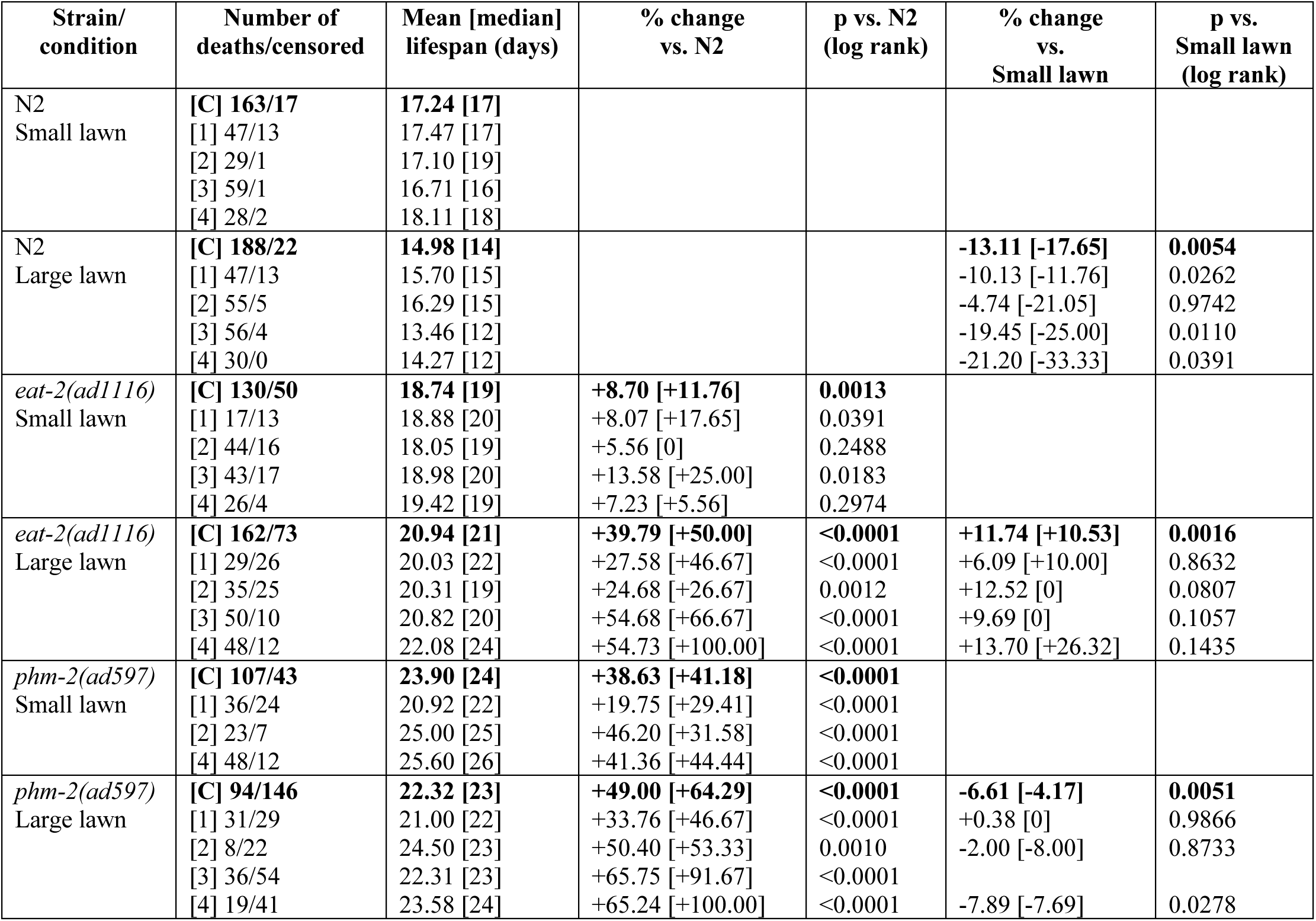
Lifespan analysis of *eat-2* and *phm-2* mutants on small and large bacterial lawns (20°C)

## References

Austad, S.N. and Kristan, D.M., 2003. Are mice calorically restricted in nature? Aging Cell. 2, 201–7.

Avery, L., 1993. The genetics of feeding in *Caenorhabditis elegans*. Genetics. 133, 897–917.

Brenner, S., 1974. The genetics of *Caenorhabditis elegans*. Genetics. 77, 71–94.

Burkewitz, K., Morantte, I., Weir, H., Robin Yeo, R., Zhang, Y., Huynh, F., Ilkayeva, O., Hirschey, M., Grant, A. and Mair, W., 2015. Neuronal CRTC-1 governs systemic mitochondrial metabolism and lifespan via a catecholamine signal. Cell. 160 842–855.

Cabreiro, F. and Gems, D., 2010. Treating aging: progress toward dietary restriction mimetics. F1000 Biol Rep. 2, 76.

Calvert, S., Tacutu, R., Sharifi, S., Teixeira, R., Ghosh, P. and de Magalhães, J.P., 2016. A network pharmacology approach reveals new candidate caloric restriction mimetics in *C. elegans*. Aging Cell. 15, 256–266.

Chang, Y.H., Chi, A.Q., Ren, Y.C., Mu, X.P., Tao, B.B., Shao, Z. and Zhu, Y.C., 2026. fln-2 isoform-specifically regulates *Caenorhabditis elegans* health span by affecting pharyngeal function. Sci Rep. 6, 8363.

Chow, D.K., Glenn, C.F., Johnston, J.L., Goldberg, I.G. and Wolkow, C.A., 2006. Sarcopenia in the *Caenorhabditis elegans* pharynx correlates with muscle contraction rate over lifespan. Exp Gerontol. 41, 252–60.

Darby, C., Hsu, J.W., Ghori, N. and Falkow, S., 2002. *Caenorhabditis elegans*: plague bacteria biofilm blocks food intake. Nature. 417, 243–4.

Di Giosia, P., Stamerra, C.A., Giorgini, P., Jamialahamdi, T., Butler, A.E. and Sahebkar, A., 2022. The role of nutrition in inflammaging. Ageing Res Rev. 77, 101596.

Garigan, D., Hsu, A.L., Fraser, A.G., Kamath, R.S., Ahringer, J. and Kenyon, C., 2002. Genetic analysis of tissue aging in *Caenorhabditis elegans*: a role for heat-shock factor and bacterial proliferation. Genetics. 161, 1101–1112.

Gems, D. and Riddle, D.L., 2000. Genetic, behavioral and environmental determinants of male longevity in *Caenorhabditis elegans*. Genetics. 154, 1597–1610.

Greer, E.L. and Brunet, A., 2009. Different dietary restriction regimens extend lifespan by both independent and overlapping genetic pathways in *C. elegans*. Aging Cell. 8, 113–27.

Hasegawa, A., Iwasaka, H., Hagiwara, S., Asai, N., Nishida, T. and Noguchi, T., 2012. Alternate day calorie restriction improves systemic inflammation in a mouse model of sepsis induced by cecal ligation and puncture. J Surg Res. 174, 136–41.

Heestand, B., Shen, Y., Liu, W., Magner, D., Storm, N., Meharg, C., Habermann, B. and Antebi, A., 2013. Dietary restriction induced longevity is mediated by nuclear receptor NHR-62 in *Caenorhabditis elegans*. PLOS Genet. 9, e1003651.

Hsiung, K.C., Chapman, H., Wei, X., Sun, X., Rawlinson, I. and Gems, D., 2026. Effects of knockdown of autophagy-specifying genes on *C. elegans* mutant longevity are highly condition dependent. eLife. In revision.

Kaeberlein, T., Smith, E., Tsuchiya, M., Welton, K., Thomas, J., Fields, S., Kennedy, B. and Kaeberlein, M., 2006. Lifespan extension in *Caenorhabditis elegans* by complete removal of food. Aging Cell. 5, 487–494.

Kapahi, P., Kaeberlein, M. and Hansen, M., 2016. Dietary restriction and lifespan: Lessons from invertebrate models. Ageing Res Rev. 39, 3–14.

Kauffman, A., Ashraf, J., Corces-Zimmerman, M., Landis, J. and Murphy, C., 2010. Insulin signaling and dietary restriction differentially influence the decline of learning and memory with age. PLOS Biol. 8, e1000372.

Kennedy, B.K. and Lamming, D.W., 2016. The Mechanistic Target of Rapamycin: The Grand ConducTOR of Metabolism and Aging. Cell Metab. 23, 990–1003.

Klass, M.R., 1977. Aging in the nematode *Caenorhabditis elegans*: major biological and environmental factors influencing life span. Mech Ageing Develop. 6, 413–429.

Kumar, S., Egan, B., Kocsisova, Z., Schneider, D., Murphy, J., Diwan, A. and Kornfeld, K., 2019. Lifespan extension in *C. elegans* caused by bacterial colonization of the intestine and subsequent activation of an innate immune response. Develop Cell. 49, 100–117.

Labrousse, A., Chauvet, S., Couillault, C., Kurz, C.L. and Ewbank, J.J., 2000. *Caenorhabditis elegans* is a model host for *Salmonella typhimurium*. Curr Biol. 10, 1543–5.

Lakowski, B. and Hekimi, S., 1998. The genetics of caloric restriction in *Caenorhabditis elegans*. Proc Natl Acad Sci USA. 95, 13091–13096.

Mattison, J.A., Colman, R.J., Beasley, T.M., Allison, D.B., Kemnitz, J.W., Roth, G.S., Ingram, D.K., Weindruch, R., de Cabo, R. and Anderson, R.M., 2017. Caloric restriction improves health and survival of rhesus monkeys. Nat Commun. 8, 14063.

Mattison, J.A., Roth, G.S., Beasley, T.M., Tilmont, E.M., Handy, A.M., Herbert, R.L., Longo, D.L., Allison, D.B., Young, J.E., Bryant, M., Barnard, D., Ward, W.F., Qi, W., Ingram, D.K. and de Cabo, R., 2012. Impact of caloric restriction on health and survival in rhesus monkeys from the NIA study. Nature. 489, 318–21.

McKay, J.P., Raizen, D.M., Gottschalk, A., Schafer, W.R. and Avery, L., 2004. *eat-2* and *eat-18* are required for nicotinic neurotransmission in the *Caenorhabditis elegans* pharynx. Genetics. 166, 161–169.

Mosser, T., Matic, I. and Leroy, M., 2011. Bacterium-induced internal egg hatching frequency is predictive of life span in *Caenorhabditis elegans* populations. Appl Environ Microbiol. 77, 8189–8192.

Most, J., Tosti, V., Redman, L.M. and Fontana, L., 2017. Calorie restriction in humans: An update. Ageing Res Rev. 39, 36–45.

Nelson, J.F., Gosden, R.G. and Felicio, L.S., 1985. Effect of dietary restriction on estrous cyclicity and follicular reserves in aging C57BL/6J mice. Biol Reprod. 32, 515–22.

Panowski, S.H., Wolff, S., Aguilaniu, H., Durieux, J. and Dillin, A., 2007. PHA-4/Foxa mediates diet-restriction-induced longevity of *C. elegans*. Nature. 447, 550–5.

Park, S.-K., Link, C. and Johnson, T., 2010. Life-span extension by dietary restriction is mediated by NLP-7 signaling and coelomocyte endocytosis in *C. elegans*. FASEB J. 24, 383–392.

Pifferi, F., Terrien, J., Marchal, J., Dal-Pan, A., Djelti, F., Hardy, I., Chahory, S., Cordonnier, N., Desquilbet, L., Hurion, M., Zahariev, A., Chery, I., Zizzari, P., Perret, M., Epelbaum, J., Blanc, S., Picq, J.L., Dhenain, M. and Aujard, F., 2018. Caloric restriction increases lifespan but affects brain integrity in grey mouse lemur primates. Commun Biol. 1, 30.

Piper, M.D. and Partridge, L., 2007. Dietary restriction in Drosophila: delayed aging or experimental artefact? PLOS Genet. 3, e57.

Podshivalova, K., Kerr, R. and Kenyon, C., 2017. How a mutation that slows aging can also disproportionately extend end-of-life decrepitude. Cell Reports. 19, 441–450.

Prinz, F., Schlange, T. and Asadullah, K., 2011. Believe it or not: how much can we rely on published data on potential drug targets? Nat Rev Drug Discov. 10, 712.

Ritchie, S., 2020. Science Fictions. Exposing Fraud, Bias, Negligence and Hype in Science, Vintage (Penguin Random House), London, UK.

Sanchez-Blanco, A. and Kim, S.K., 2011. Variable pathogenicity determines individual lifespan in *Caenorhabditis elegans*. PLOS Genet. 7, e1002047.

Schmauck-Medina, T., Lautrup, S., Di Francesco, A., Mitchell, S.J., Clemmensen, C., Partridge, L., Roth, G.S., Anderson, R.M., Mattison, J.A., de Cabo, R., Speakman, J.R., Richardson, A., Ingram, D.K., Weindruch, R., Mattson, M.P. and Fang, E.F., 2026. Dietary restriction in aging and longevity. Nat Aging. 6, 485–505.

Singh, J. and Aballay, A., 2019. Microbial colonization activates an immune fight-and-flight response via neuroendocrine signaling. Develop Cell. 49, 89–99.

Statzer, C., Reichert, P., Dual, J. and Ewald, C.Y., 2022. Longevity interventions temporally scale healthspan in *Caenorhabditis elegans*. iScience. 25, 103983.

Voelkl, B., Altman, N.S., Forsman, A., Forstmeier, W., Gurevitch, J., Jaric, I., Karp, N.A., Kas, M.J., Schielzeth, H., Van de Casteele, T. and Würbel, H., 2020. Reproducibility of animal research in light of biological variation. Nat Rev Neurosci. 21, 384–393.

Walker, G., Houthoofd, K., Vanfleteren, J. and Gems, D., 2005. Dietary restriction in *C. elegans*: From rate-of-living effects to nutrient sensing pathways. Exp Gerontol. 126, 929–937.

Weindruch, R., Walford, R., Fligiel, S. and Guthrie, D., 1986. The retardation of aging in mice by dietary restriction: longevity, cancer, immunity and lifetime energy intake. J Nutr. 116, 641–654.

Wu, Z., Isik, M., Moroz, N., Steinbaugh, M.J., Zhang, P. and Blackwell, T.K., 2019. Dietary restriction extends lifespan through metabolic regulation of innate immunity. Cell Metabolism. 29, 1192–1205.

Zhao, Y., Gilliat, A.F., Ziehm, M., Turmaine, M., Wang, H., Ezcurra, M., Yang, C., Phillips, G., McBay, D., Zhang, W.B., Partridge, L., Pincus, Z. and Gems, D., 2017. Two forms of death in aging *Caenorhabditis elegans*. Nat Commun. 8, 15458.

Zhao, Y., Wang, H., Poole, R.J. and Gems, D., 2019. A *fln-2* mutation affects lethal pathology and lifespan in *C. elegans*. Nat Commun. 10, 5087.

Admasu, T.D., Batchu, K.C., Ng, L.F., Cazenave-Gassiot, A., Wenk, M.R. and Gruber, J., 2018. Lipid profiling of *C. elegans* strains administered pro-longevity drugs and drug combinations. Sci Data. 5, 180231.

Artal-Sanz, M. and Tavernarakis, N., 2009. Prohibitin couples diapause signalling to mitochondrial metabolism during ageing in *C. elegans*. Nature. 461, 793–7.

Bansal, A., Zhu, L.J., Yen, K. and Tissenbaum, H.A., 2015. Uncoupling lifespan and healthspan in *Caenorhabditis elegans* longevity mutants. Proc Natl Acad Sci U S A. 112, E277–86.

Campos, J.C., Marchesi Bozi, L.H., Krum, B., Grassmann Bechara, L.R., Ferreira, N.D., Arini, G.S., Albuquerque, R.P., Traa, A., Ogawa, T., van der Bliek, A.M., Beheshti, A., Chouchani, E.T., Van Raamsdonk, J.M., Blackwell, T.K. and Ferreira, J.C.B., 2023. Exercise preserves physical fitness during aging through AMPK and mitochondrial dynamics. Proc Natl Acad Sci U S A. 120, e2204750120.

Carrano, A., Liu, Z., Dillin, A. and Hunter, T., 2009. A conserved ubiquitination pathway determines longevity in response to diet restriction. Nature. 460, 396–399.

Chamoli, M., Singh, A., Malik, Y. and Mukhopadhyay, A., 2014. A novel kinase regulates dietary restriction-mediated longevity in *Caenorhabditis elegans*. Aging Cell. 13, 641–655.

Chen, D., Thomas, E. and Kapahi, P., 2009. HIF-1 modulates dietary restriction-mediated lifespan extension via IRE-1 in *Caenorhabditis elegans*. PLoS Genet. 5, e1000486.

Chen, W., Lin, H.R., Wei, C.M., Luo, X.H., Sun, M.L., Yang, Z.Z., Chen, X.Y. and Wang, H.B., 2018. Echinacoside, a phenylethanoid glycoside from Cistanche deserticola, extends lifespan of *Caenorhabditis elegans* and protects from Aβ-induced toxicity. Biogerontology. 19, 47–65.

Chen, Y., Onken, B., Chen, H., Zhang, X., Driscoll, M., Cao, Y. and Huang, Q., 2020. Healthy lifespan extension mediated by oenothein B isolated from *Eucalyptus grandis* × *Eucalyptus urophylla* GL9 in *Caenorhabditis elegans*. Food Funct. 11, 2439–2450.

Ching, T.T., Paal, A.B., Mehta, A., Zhong, L. and Hsu, A.L., 2010. *drr-2* encodes an eIF4H that acts downstream of TOR in diet-restriction-induced longevity of *C. elegans*. Aging Cell. 9, 545–57.

Cho, J. and Park, Y., 2023. Kahweol, a coffee diterpene, increases lifespan via insulin/insulin-like growth factor-1 and AMP-activated protein kinase signaling pathways in *Caenorhabditis elegans*. Curr Res Food Sci. 7, 100618.

Cornwell, A.B., Zhang, Y., Thondamal, M., Johnson, D.W., Thakar, J. and Samuelson, A.V., 2024. The *C. elegans* Myc-family of transcription factors coordinate a dynamic adaptive response to dietary restriction. Geroscience. 46, 4827–4854.

Costa, M.D., Da Silva, J.D., Almeida, D., Pereira-Sousa, J., Vilasboas-Campos, D., Fernandes, J.H., Teixeira-Castro, A. and Maciel, P., 2025. Differential effects of lifespan-extending genetic manipulations in an animal model of MJD/SCA3. Mech Ageing Dev. 225, 112064.

Crawford, D., Libina, N. and Kenyon, C., 2007. *Caenorhabditis elegans* integrates food and reproductive signals in lifespan determination. Aging Cell. 6, 715–21.

Curtis, R., O’Connor, G. and DiStefano, P.S., 2006. Aging networks in *Caenorhabditis elegans*: AMP-activated protein kinase (aak-2) links multiple aging and metabolism pathways. Aging Cell. 5, 119–26.

Dawson, Z.D., Sundaramoorthi, H., Regmi, S., Zhang, B., Morrison, S., Fielder, S.M., Zhang, J.R., Hoang, H., Perlmutter, D.H., Luke, C.J., Silverman, G.A. and Pak, S.C., 2024. A fluorescent reporter for rapid assessment of autophagic flux reveals unique autophagy signatures during *C. elegans* post-embryonic development and identifies compounds that modulate autophagy. Autophagy Rep. 3, 2371736.

Ding, S.S., Romenskyy, M., Sarkisyan, K.S. and Brown, A.E.X., 2020. Measuring *Caenorhabditis elegans* spatial foraging and food intake using bioluminescent bacteria. Genetics. 214, 577–587.

Essmann, C.L., Martinez-Martinez, D., Pryor, R., Leung, K.-Y., Krishnan, K.B., Lui, P.P., Greene, N.D.E., Brown, A.E.X., Pawar, V.M., Srinivasan, M.A. and Cabreiro, F., 2020. Mechanical properties measured by atomic force microscopy define health biomarkers in ageing *C. elegans*. Nat Commun. 11, 1043.

Farias-Pereira, R., Park, C.S. and Park, Y., 2020. Kahweol Reduces Food Intake of *Caenorhabditis elegans*. J Agric Food Chem. 68, 9683–9689.

Galbadage, T. and Hartman, P.S., 2008. Repeated temperature fluctuation extends the life span of *Caenorhabditis elegans* in a *daf-16*-dependent fashion. Mech Ageing Dev. 129, 507–14.

Gao, A.W., Smith, R.L., van Weeghel, M., Kamble, R., Janssens, G.E. and Houtkooper, R.H., 2018. Identification of key pathways and metabolic fingerprints of longevity in *C. elegans*. Exp Gerontol. 113, 128–140.

García-Casas, P., Arias-Del-Val, J., Alvarez-Illera, P., Wojnicz, A., de Los Ríos, C., Fonteriz, R.I., Montero, M. and Alvarez, J., 2018. The Neuroprotector Benzothiazepine CGP37157 Extends Lifespan in *C. elegans* Worms. Front Aging Neurosci. 10, 440.

Gelino, S., Chang, J., Kumsta, C., She, X., Davis, A., Nguyen, C., Panowski, S. and Hansen, M., 2016. Intestinal autophagy improves healthspan and longevity in *C. elegans* during dietary restriction. PLoS Genet. 12, e1006135.

Govindhan, T., Amirthalingam, M., Govindan, S., Duraisamy, K., Cho, J.H., Tawata, S., Periyakali, S.B. and Palanisamy, S., 2024. Diosgenin intervention: targeting lipophagy to counter high glucose diet-induced lipid accumulation and lifespan reduction. 3 Biotech. 14, 171.

Hahm, J.H., Jeong, C. and Nam, H.G., 2019. Diet restriction-induced healthy aging is mediated through the immune signaling component ZIP-2 in *Caenorhabditis elegans*. Aging Cell. 18, e12982.

Hansen, M., Chandra, A., Mitic, L.L., Onken, B., Driscoll, M. and Kenyon, C., 2008. A role for autophagy in the extension of lifespan by dietary restriction in *C. elegans*. PLoS Genet. 4, e24.

Hansen, M., Taubert, S., Crawford, D., Libina, L., Lee, S.-J. and Kenyon, C., 2007. Lifespan extension by conditions that inhibit translation in *Caenorhabditis elegans*. Aging Cell. 6, 95–110.

Hsu, A., Murphy, C. and Kenyon, C., 2003. Regulation of aging and age-related disease by DAF-16 and heat-shock factor. Science. 300, 1142–1145.

Huang, C., Xiong, C. and Kornfeld, K., 2004. Measurements of age-related changes of physiological processes that predict lifespan of *Caenorhabditis elegans*. Proc Natl Acad Sci U S A. 101, 8084–8089.

Iser, W. and Wolkow, C., 2007. DAF-2/insulin-like signaling in C. elegans modifies effects of dietary restriction and nutrient stress on aging, stress and growth. PLoS One. 2, e1240.

Jia, K. and Levine, B., 2007. Autophagy is required for dietary restriction-mediated life span extension in *C. elegans*. Autophagy. 3, 597–599.

Jiang, S., Jiang, C.P., Cao, P., Liu, Y.H., Gao, C.H. and Yi, X.X., 2022. Sonneradon A Extends Lifespan of *Caenorhabditis elegans* by Modulating Mitochondrial and IIS Signaling Pathways. Mar Drugs. 20.

Juozaityte, V., Pregnolato, C., Abay-Nørgaard, S., Rausch, D.M., McIntyre, R.L., Gerhart-Hines, Z., Pers, T.H., Salcini, A.E. and Clemmensen, C., 2026. The NMDA Receptor Antagonist Memantine Modulates Aging and Stress Resilience. Aging Cell. 25, e70303.

Kim, S.H., Kim, B.K. and Park, S.K., 2018. Selenocysteine mimics the effect of dietary restriction on lifespan via SKN-1 and retards age-associated pathophysiological changes in *Caenorhabditis elegans*. Mol Med Rep. 18, 5389–5398.

Kozlova, A.A., Lotfi, M. and Okkema, P.G., 2019. Cross Talk with the GAR-3 Receptor Contributes to Feeding Defects in *Caenorhabditis elegans eat-2* Mutants. Genetics. 212, 231–243.

Lakowski, B. and Hekimi, S., 1998. The genetics of caloric restriction in *Caenorhabditis elegans*. Proc Natl Acad Sci U S A. 95, 13091–13096.

Lee, S.J., Hwang, A.B. and Kenyon, C., 2010. Inhibition of respiration extends *C. elegans* life span via reactive oxygen species that increase HIF-1 activity. Curr Biol. 20, 2131–6.

Lin, Z.H., Chang, S.Y., Shen, W.C., Lin, Y.H., Shen, C.L., Liao, S.B., Liu, Y.C., Chen, C.S., Ching, T.T. and Wang, H.D., 2022. Isocitrate Dehydrogenase Alpha-1 Modulates Lifespan and Oxidative Stress Tolerance in *Caenorhabditis elegans*. Int J Mol Sci. 24, 612.

Lu, M., Tan, L., Zhou, X.G., Yang, Z.L., Zhu, Q., Chen, J.N., Luo, H.R. and Wu, G.S., 2020a. Secoisolariciresinol Diglucoside Delays the Progression of Aging-Related Diseases and Extends the Lifespan of *Caenorhabditis elegans* via DAF-16 and HSF-1. Oxid Med Cell Longev. 2020, 1293935.

Lu, M., Tan, L., Zhou, X.G., Yang, Z.L., Zhu, Q., Chen, J.N., Luo, H.R. and Wu, G.S., 2020b. Tectochrysin increases stress resistance and extends the lifespan of *Caenorhabditis elegans* via FOXO/DAF-16. Biogerontology. 21, 669–682.

Mehta, R., Steinkraus, K., Sutphin, G., Ramos, F., Shamieh, L., Huh, A., Davis, C., Chandler-Brown, D. and Kaeberlein, M., 2009. Proteasomal regulation of the hypoxic response modulates aging in *C. elegans*. Science. 324, 1196–1198.

Meng, F., Li, J., Rao, Y., Wang, W. and Fu, Y., 2018. Gengnianchun Extends the Lifespan of *Caenorhabditis elegans* via the Insulin/IGF-1 Signalling Pathway. Oxid Med Cell Longev. 2018, 4740739.

Mi, X.N., Wang, L.F., Hu, Y., Pan, J.P., Xin, Y.R., Wang, J.H., Geng, H.J., Hu, S.H., Gao, Q. and Luo, H.M., 2018. Methyl 3,4-Dihydroxybenzoate Enhances Resistance to Oxidative Stressors and Lifespan in *C. elegans* Partially via *daf-2*/*daf-16*. Int J Mol Sci. 19, 1670.

Mir, D.A., Cox, M., Horrocks, J., Ma, Z. and Rogers, A., 2024. Roles of Progranulin and FRamides in Neural Versus Non-Neural Tissues on Dietary Restriction-Related Longevity and Proteostasis in *C. elegans*. J Clin Med Sci. 8, 276.

Murakami, H. and Murakami, S., 2007. Serotonin receptors antagonistically modulate *Caenorhabditis elegans* longevity. Aging Cell. 6, 483–8.

Ng, L.T., Ng, L.F., Tang, R.M.Y., Barardo, D., Halliwell, B., Moore, P.K. and Gruber, J., 2020. Lifespan and healthspan benefits of exogenous H_2_S in *C. elegans* are independent from effects downstream of *eat-2* mutation. NPJ Aging Mech Dis. 6, 6.

Onken, B. and Driscoll, M., 2010. Metformin induces a dietary restriction-like state and the oxidative stress response to extend *C. elegans* Healthspan via AMPK, LKB1, and SKN-1. PLoS One. 5, e8758.

Palikaras, K., Lionaki, E. and Tavernarakis, N., 2015. Coordination of mitophagy and mitochondrial biogenesis during ageing in *C. elegans*. Nature. 521, 525–8.

Pan, K.Z., Palter, J.E., Rogers, A.N., Olsen, A., Chen, D., Lithgow, G.J. and Kapahi, P., 2007. Inhibition of mRNA translation extends lifespan in *Caenorhabditis elegans*. Aging Cell. 6, 111–9.

Petrascheck, M., Ye, X. and Buck, L.B., 2007. An antidepressant that extends lifespan in adult *Caenorhabditis elegans*. Nature. 450, 553–6.

Pinkston, J.M., Garigan, D., Hansen, M. and Kenyon, C., 2006. Mutations that increase the life span of *C. elegans* inhibit tumor growth. Science. 313, 971–5.

Powolny, A.A., Singh, S.V., Melov, S., Hubbard, A. and Fisher, A.L., 2011. The garlic constituent diallyl trisulfide increases the lifespan of *C. elegans* via *skn-1* activation. Exp Gerontol. 46, 441–52.

Schlotterer, A., Kukudov, G., Bozorgmehr, F., Hutter, H., Du, X., Oikonomou, D., Ibrahim, Y., Pfisterer, F., Rabbani, N., Thornalley, P., Sayed, A., Fleming, T., Humpert, P., Schwenger, V., Zeier, M., Hamann, A., Stern, D., Brownlee, M., Bierhaus, A., Nawroth, P. and Morcos, M., 2009. *C. elegans* as model for the study of high glucose–mediated life span reduction. Diabetes. 58, 2450–2456.

Shpigel, N., Shemesh, N., Kishner, M. and Ben-Zvi, A., 2019. Dietary restriction and gonadal signaling differentially regulate post-development quality control functions in *Caenorhabditis elegans*. Aging Cell. 18, e12891.

Singh, A., Kumar, N., Matai, L., Jain, V., Garg, A. and Mukhopadhyay, A., 2016. A chromatin modifier integrates insulin/IGF-1 signalling and dietary restriction to regulate longevity. Aging Cell. 15, 694–705.

Staab, T.A., McIntyre, G., Wang, L., Radeny, J., Bettcher, L., Guillen, M., Peck, M.P., Kalil, A.P., Bromley, S.P., Raftery, D. and Chan, J.P., 2023. The lipidomes of *C. elegans* with mutations in asm-3/acid sphingomyelinase and hyl-2/ceramide synthase show distinct lipid profiles during aging. Aging (Albany NY). 15, 650–674.

Sun, Y., Li, M., Zhao, D., Li, X., Yang, C. and Wang, X., 2020. Lysosome activity is modulated by multiple longevity pathways and is important for lifespan extension in *C. elegans*. eLife. 9, e55745.

Syntichaki, P., Troulinaki, K. and Tavernarakis, N., 2007. eIF4E function in somatic cells modulates ageing in *Caenorhabditis elegans*. Nature. 445, 922–6.

Tabrez, S., Sharma, R., Jain, V., Siddiqui, A. and Mukhopadhyay, A., 2017. Differential alternative splicing coupled to nonsense-mediated decay of mRNA ensures dietary restriction-induced longevity. Nat Commun. 8, 306.

Toth, M.L., Sigmond, T., Borsos, E., Barna, J., Erdelyi, P., Takacs-Vellai, K., Orosz, L., Kovacs, A.L., Csikos, G., Sass, M. and Vellai, T., 2008. Longevity pathways converge on autophagy genes to regulate life span in *Caenorhabditis elegans*. Autophagy. 4, 330–8.

Van Raamsdonk, J.M. and Hekimi, S., 2009. Deletion of the mitochondrial superoxide dismutase *sod-2* extends lifespan in *Caenorhabditis elegans*. PLOS Genet. 5, e1000361.

Vilchez, D., Morantte, I., Liu, Z., Douglas, P., Merkwirth, C., Rodrigues, A., Manning, G. and Dillin, A., 2012. RPN-6 determines *C. elegans* longevity under proteotoxic stress conditions. Nature. 489, 263–268.

Viri, V., Cornaglia, M., Atakan, H.B., Lehnert, T. and Gijs, M.A.M., 2020. An in vivo microfluidic study of bacterial transit in *C. elegans* nematodes. Lab Chip. 20, 2696–2708.

Wang, X., Li, H., Liu, Y., Wu, H., Wang, H., Jin, S., Lu, Y., Chang, S., Liu, R., Peng, Y., Guo, Z. and Wang, X., 2020. Velvet antler methanol extracts (MEs) protects against oxidative stress in *Caenorhabditis elegans* by SKN-1. Biomed Pharmacother. 121, 109668.

Wang, Y. and Tissenbaum, H.A., 2006. Overlapping and distinct functions for a *Caenorhabditis elegans* SIR2 and DAF-16/FOXO. Mech Ageing Dev. 127, 48–56.

Yang, Z.Z., Yu, Y.T., Lin, H.R., Liao, D.C., Cui, X.H. and Wang, H.B., 2018. *Lonicera japonica* extends lifespan and healthspan in *Caenorhabditis elegans*. Free Radic Biol Med. 129, 310–322.

Yee, C., Yang, W. and Hekimi, S., 2014. The intrinsic apoptosis pathway mediates the pro-longevity response to mitochondrial ROS in C. elegans. Cell. 157, 897–909.

Yuan, Y., Kadiyala, C.S., Ching, T.T., Hakimi, P., Saha, S., Xu, H., Yuan, C., Mullangi, V., Wang, L., Fivenson, E., Hanson, R.W., Ewing, R., Hsu, A.L., Miyagi, M. and Feng, Z., 2012. Enhanced energy metabolism contributes to the extended life span of calorie-restricted *Caenorhabditis elegans*. J Biol Chem. 287, 31414–26.

Zaarur, N., Desevin, K., Mackenzie, J., Lord, A., Grishok, A. and Kandror, K.V., 2019. ATGL-1 mediates the effect of dietary restriction and the insulin/IGF-1 signaling pathway on longevity in *C. elegans*. Mol Metab. 27, 75–82.

Zeng, W.Y., Tan, L., Han, C., Zheng, Z.Y., Wu, G.S., Luo, H.R. and Li, S.L., 2021. Trigonelline Extends the Lifespan of *C. Elegans* and Delays the Progression of Age-Related Diseases by Activating AMPK, DAF-16, and HSF-1. Oxid Med Cell Longev. 2021, 7656834.

Zhu, B., Jo, K., Yang, P., Tohti, J., Fei, J. and Abudukerim, K., 2019. Aiweixin, a Traditional Uyghur Medicinal Formula, Extends the Lifespan of *Caenorhabditis elegans*. Evid Based Complement Alternat Med. 2019, 3684601.

